# Bypass of Blocking Lesions by RNAPII Impairs the Transcriptional DNA Damage Response

**DOI:** 10.1101/2025.04.28.651040

**Authors:** Carolina P. Bañuelos, Lucas D. Caeiro, Pradeepkumar R. Cingaram, Felipe Beckedorff, Lluis Morey, Daniel Bilbao Cortes, Ramin Shiekhattar, Ramiro E. Verdun

## Abstract

Ultraviolet (UV) irradiation and platinum-based drugs generate bulky DNA lesions that impede transcription elongation by RNA Polymerase II (RNAPII). This transcriptional block triggers a coordinated stress response involving transcription-coupled nucleotide excision repair (TC-NER), removal and degradation of the stalled RNAPII, and global transcriptional shutdown. However, the molecular and cellular consequences of RNAPII bypassing such lesions remain unclear.

Here, we identify the acetyltransferase p300 as a key regulator of this transcriptional stress response. p300 interacts with stalled RNAPII and promotes its removal and degradation through a USP7-dependent mechanism. Remarkably, in p300-deficient cells, RNAPII bypasses DNA lesions, allowing transcription to persist despite DNA damage and leading to the production of full-length mRNAs. This sustained transcriptional activity without DNA lesion repair results in increased genome instability and reduced cellular proliferation capacity. These findings reveal the biological consequences of transcribing through transcription-blocking lesions.

## Main Text

Gene transcription is crucial for maintaining cell identity and survival. However, DNA damage, particularly from bulky lesions such as cyclobutane pyrimidine dimers (CPDs) and pyrimidine-pyrimidone (6–4) photoproducts (6–4PPs) caused by UV radiation, severely hinders this process (Nieto Moreno et al., 2023). These lesions can block RNA Polymerase II (RNAPII) elongation (Agapov et al., 2022; Brueckner et al., 2007; Wang et al., 2018) - a problem also observed when cells are exposed to platinum-based chemotherapy drugs like oxaliplatin and cisplatin, used in treating numerous cancer patients (Kelland, 2007; Zhang et al., 2022).

To address this, cells employ the transcription-coupled nucleotide excision repair (TC-NER) pathway, which specifically removes bulky DNA lesions from actively transcribed DNA strands (Gregersen and Svejstrup, 2018; Lans et al., 2019). Defects in TC-NER genes are linked to disorders such as Cockayne syndrome (CS) and UV-sensitive syndrome (UV^S^S) (Henning et al., 1995; Nakazawa et al., 2012; Troelstra et al., 1992; Zhang et al., 2012). The process begins when the CSB protein, part of the TC-NER complex, detects and binds to a stalled RNAPII, attempting to resume transcription by pushing RNAPII past the lesion (Kokic et al., 2021; Li et al., 2014; van den Boom et al., 2004; van Gool et al., 1997; Xu et al., 2017).

If unsuccessful, this leads to the recruitment of additional repair proteins, including CSA, DDB1, CUL4A, and RBX1, which form the CRL4^CSA^ complex (Kokic et al., 2024; Liebelt et al., 2020; Nakazawa et al., 2020; Tufegdzic Vidakovic et al., 2020; van der Weegen et al., 2020). This complex ubiquitylates RPB1, the biggest subunit of RNAPII, signaling for its removal from DNA and subsequent degradation (Anindya et al., 2007; Bregman et al., 1996; Kleiman et al., 2005; Nakazawa et al., 2020; Ratner et al., 1998; Starita et al., 2005; Tufegdzic Vidakovic et al., 2020; Yasukawa et al., 2008). The extent of this damage and the subsequent degradation of RNAPII can drastically reduce the available RNAPII pool, necessitating a complete shutdown of transcription initiation across the cell (Nakazawa et al., 2020; Tufegdzic Vidakovic et al., 2020). This shutdown is vital for DNA repair and the quick resumption of normal transcription functions, which are crucial for cell survival.

Previous research has extensively explored the mechanisms behind transcriptional shutdown and recovery after DNA damage (Adam et al., 2013; Epanchintsev et al., 2017; Gregersen and Svejstrup, 2018; Lans et al., 2019; Noe Gonzalez et al., 2021; Vermeulen and Fousteri, 2013). Furthermore, amino acid substitutions in the largest subunit of yeast RNAPII—known to enhance processivity and reduce fidelity—have been shown to increase the bypass of blocking lesions, impair TC-NER–mediated repair, and affect cell proliferation (Li et al., 2014; Walmacq et al., 2012). Yet, the consequences of RNAPII bypassing transcription-blocking lesions and sustaining transcription through damaged chromatin in mammalian cells remain unclear. Our study reveals that inhibition of the E1A binding protein p300 allows RNAPII to bypass these lesions. This bypass results in continuous transcription, increased DNA damage and reduced cell proliferation capacity. This novel stress triggered by RNAPII bypassing blocking lesions offers new insights into resistance mechanisms against DNA damage.

### P300 is required for transcriptional shutdown post DNA damage

After a global shutdown of transcription due to RNAPII stalling at blocking DNA lesions, it is imperative for cells to restore transcription activity for survival. Although several factors have been associated with this recovery of transcription activity (Adam et al., 2013; Epanchintsev et al., 2017; Proietti-De-Santis et al., 2006; Vermeulen and Fousteri, 2013), the underlying mechanism is still not fully understood. To gain further insights into this process, we initiated our study by investigating the potential involvement of the acetyltransferase p300, a well-characterized co-activator of transcription, in the resumption of transcription following DNA damage. Using 5-ethynyl uridine (5-EU) to track nascent transcript levels, we compared responses in wild-type (WT) and p300-deficient human cells, both before and after DNA damage. Post UV irradiation, WT cells exhibited the expected decrease in transcription activity at two hours, followed by a return to normal levels after 24 hours (**Figures 1A-1C**). Contrastingly, p300-deficient cells, through either CRISPR inactivation (Δp300) or siRNA (sip300), maintained transcription levels similar to pre-damage cells, without the expected reduction in 5-EU intake (**Figures 1B and 1C**). A similar trend was observed upon inhibiting p300 acetyltransferase activity with the small molecule A-485 (Lasko et al., 2017), which affected the acetylation of p300 targets but not cell cycle progression or proliferation (**Figures 1C and S1A-S1D**). This effect was seen in HeLa (cervical cancer), IMR90 (primary lung fibroblasts), and U2OS (osteosarcoma) cells post exposure to UV (**Figures 1C and 1D**). These studies suggested that p300 plays a role on the transcriptional shutdown following DNA damage that produces transcription-blocking lesions.

**Figure 1.**
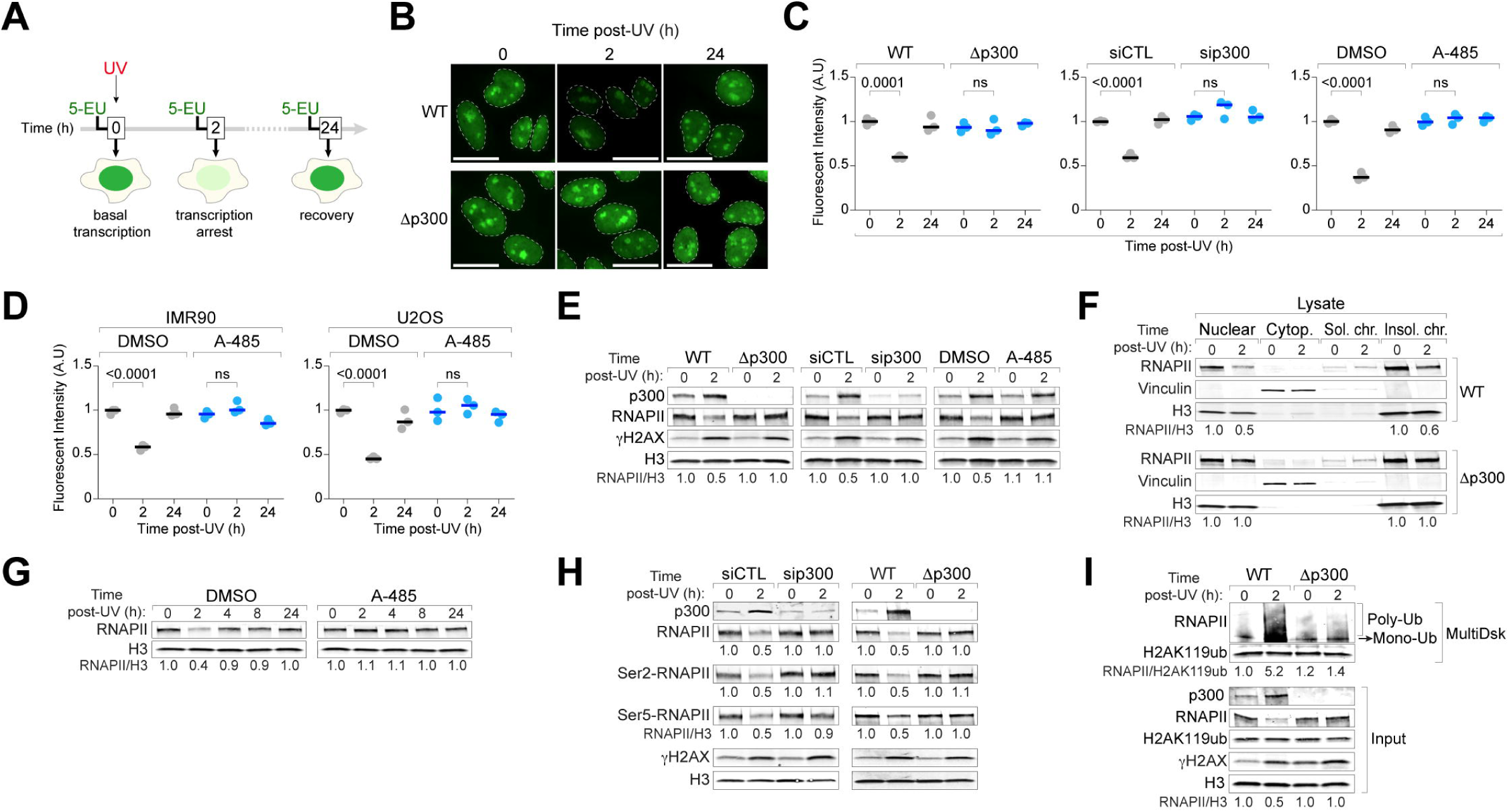
P300 is required for RNAPII degradation post DNA damage. **(A)** Schematic illustrating the strategy for the labeling and visualization of nascent RNA transcripts using 5-Ethynyl uridine (5-EU) before or after UV-irradiation (5 J/m^2^). 5-EU was added 10 minutes before collection of each sample. **(B)** Representative fluorescence micrographs displaying the 5-EU signal at different time points following UV-exposure (5 J/m^2^). White bars = 10 μm. **(C)** Quantification of 5-EU signal in WT or Δp300 HeLa cells at the indicated time points. Each dot represents the average 5-EU signal per cell, calculated from at least one hundred cells in each experimental replicate (n=3). Samples were normalized using the signal from undamaged cells. 1 μM A-485 was added 1 h before UV-irradiation (5 J/m^2^), while siRNAs were transfected 24 h prior. Black lines indicate median values. P values from one-way ANOVA are provided. ns = not significant. **(D)** As in *C*, but using U2OS and IMR90 cells exposed to DMSO or 1 μM A-485. At least 100 cells were evaluated per sample in each experimental replicate (n=3). Black lines indicate median values. P values from one-way ANOVA are provided. ns = not significant. **(E)** Western blot (WB) analyses for the indicated proteins were performed on chromatin-enriched extracts from HeLa cells (WT, Δp300, siCTL, sip300, DMSO, or A-485) collected 2 h post UV-exposure (20 J/m^2^). A-485 (1 μM) or DMSO were added 1 h before UV-irradiation, while siRNAs were transfected 24 h prior. The ratios of RNAPII and H3 signals relative to WT, siCTL, or DMSO samples before UV damage are shown. **(F)** Cellular fractionation combined with WB analyses with the indicated antibodies and using WT or Δp300 HeLa cells. The ratios of RNAPII and H3 signals relative to samples before UV damage are shown. **(G)** WB showing RNAPII levels in HeLa cells using antibodies against RPB1 at different time points post UV-irradiation (20 J/m^2^). A-485 (1 μM) or DMSO were added 1 h before UV. The ratios of RNAPII and H3 signals relative to samples before UV damage are shown. **(H)** WB analyses for different phosphorylated forms of RNAPII at 0 and 2 h post UV-irradiation. **(I)** WB from MultiDSK pull-down assays with WT or Δp300 WT or Δp300 HeLa cells at 0 and 2 h post UV-irradiation. All experiments were performed in at least three independent experimental replicates.

In mammalian cells, DNA damage-induced transcriptional elongation and initiation shutdown are linked to RNAPII stalling and proteosome-mediated degradation respectively (**Figure S1E**) (Anindya et al., 2007; Bregman et al., 1996; Kleiman et al., 2005; Nakazawa et al., 2020; Ratner et al., 1998; Starita et al., 2005; Tufegdzic Vidakovic et al., 2020; Yasukawa et al., 2008). Based on our previous results, we next investigated if p300 deficiency impacts RNAPII stability after DNA damage. For these studies, we utilized RPB1 (POLR2A), the catalytic subunit of RNAPII, as a marker to determine its protein levels. Unlike in HeLa control cells, RNAPII levels did not decrease after UV damage in p300-deficient cells, maintaining consistent protein levels in the nuclear and chromatin fractions at different UV doses (**Figures 1E, 1F and S1F**). This was corroborated in U2OS and IMR90 cells treated with A-485 and subjected to UV exposure (**Figure S1G**). However, cells deficient in CREB-binding protein (CREBBP), a homolog of p300, showed a reduction in RNAPII levels after DNA damage (**Figure S1H**), demonstrating that this phenotype is specific to p300 activity. Importantly, time-course experiments revealed that the effect of p300 inhibition on the removal of RNAPII from damaged chromatin was not due to a delay in the response (**Figure 1G**). Additionally, p300 inhibition did not affect the phosphorylation of the C-terminal domain of RNAPII at serine 2 (Ser2-RNAPII, which is enriched during elongation) or at serine 5 (Ser5-RNAPII, which is enriched at transcription start sites, or TSS) (**Figure 1H**). These results suggest that RNAPII activity was not altered after DNA damage in p300-deficient cells.

Stalled RNAPII degradation post DNA damage relies on RPB1 poly-ubiquitylation by the E3 ligase complex CRL4^CSA^ (Kokic et al., 2024; Liebelt et al., 2020; Nakazawa et al., 2020; Tufegdzic Vidakovic et al., 2020; van der Weegen et al., 2020). To evaluate this, we conducted MultiDsk pulldown assays (Wilson et al., 2012) to measure the ubiquitylation levels of RPB1 before and after DNA damage in WT and Δp300 cells. We used histone H2A ubiquitylated at lysine 119 as a control for pulldown efficiency. In contrast to WT cells, which exhibited increased poly-ubiquitylation of RPB1 upon UV exposure, p300-deficient cells displayed low levels of poly-ubiquitylated RPB1 under the same conditions (**Figure 1I**). This reduced RPB1 ubiquitylation aligns with the unaltered RNAPII protein levels and lack of transcriptional shutdown following DNA damage observed in p300-deficient cells. In addition, performing these studies in the presence of cycloheximide to inhibit *de novo* translation revealed that the effect observed in p300-deficient cells post DNA damage is caused by RNAPII degradation rather than differences in its synthesis (**Figure S1I**). These findings demonstrate a pivotal and unexpected role for p300 in RNAPII ubiquitylation, removal from damaged chromatin, and degradation, as well as in the transcriptional shutdown following DNA damage induced by UV irradiation that produce transcription blocking lesions.

### TC-NER prevents RNAPII displacement from the damaged chromatin in p300 deficient cells

CSA, the substrate-recognition factor of the DDB1–CUL4A–RBX1 ubiquitin ligase complex (CRL4^CSA^), binds to and ubiquitylates the RNAPII-CSB complex stalled at transcription-blocking lesions (TBL) (Groisman et al., 2006; Kokic et al., 2024; Liebelt et al., 2020; Nakazawa et al., 2020; Tufegdzic Vidakovic et al., 2020; van der Weegen et al., 2020). Following this, UVSSA is recruited through interaction with the ubiquitylated RNAPII, CSA, and ELOF1, subsequently bringing TFIIH into the process (Kokic et al., 2024; Liebelt et al., 2020; Nakazawa et al., 2020). Interestingly, UVSSA also recruits the ubiquitin protease USP7, which deubiquitylates CSB (Fei and Chen, 2012; Higa et al., 2018; Schwertman et al., 2012; Zhang et al., 2012). It is believed that USP7 slows down the ubiquitylation process of CSB by CRL4^CSA^ and delays its removal from the damaged chromatin. However, how the release of USP7 from the TC-NER complex is regulated to facilitate RNAPII removal from the damaged chromatin remains unclear.

The observed low ubiquitylation and steady levels of RNAPII after DNA damage in p300-deficient cells suggest a defect in its removal by TC-NER from the damaged chromatin. Also, it was previously shown that lack of ubiquitylation of the stalled RNAPII results in longer association of RNAPII and CSB with the damaged chromatin (Tufegdzic Vidakovic et al., 2020). To investigate this, we conducted immunoprecipitation (IP) assays for the elongating form of RNAPII (Ser2-RNAPII), using chromatin-enriched extracts from WT and Δp300 HeLa cells exposed or not to UV irradiation. We found that p300 accumulates at the chromatin and associates with Ser2-RNAPII after DNA damage (**Figures 2A and 2B**). Notably, p300 inhibition significantly enhanced the early association of USP7, and UVSSA with Ser2-RNAPII and damaged chromatin, while not affecting CSB, CSA, p62 (TFIIH) or CUL4A (**Figures 2A-2C, S2A and S2B**). These observations suggest a direct involvement of p300 in TC-NER and the removal of the stalled RNAPII from the chromatin. Supporting this, loss of CSB, CSA or UVSSA-but not p62 or CUL4A-inhibited the accumulation of p300 in chromatin following DNA damage (**Figures 2D and S2C**). Given that UVSSA is essential for USP7 (Schwertman et al., 2012) and p300 accumulation on damaged chromatin, these data suggest that p300 functionally interacts with UVSSA and USP7 during TC-NER. To determine whether TC-NER and USP7 contribute to the defective RNAPII removal from the damaged chromatin observed in p300-deficient cells, we inhibited TC-NER via CSB knockdown. This abolished the impact of p300 deficiency on RNAPII association with the damaged chromatin (**Figure 2E**). Similar results were observed after inhibition of USP7 with the small molecule FT671 (Turnbull et al., 2017) and in USP7 knockout (ΔUSP7) cells (**Figures 2F and S2D**). These findings demonstrate that in p300-deficient cells, TC-NER and USP7 prevent RNAPII removal from the damaged chromatin after exposure to transcription-blocking agents. Since p300 inhibition increases UVSSA and USP7 association with Ser2-RNAPII in a manner dependent on USP7 activity, these findings suggest a direct role for p300 in disengaging USP7 from the stalled RNAPII/TC-NER complex. This action could be crucial for facilitating the CRL4^CSA^-dependent polyubiquitylation and subsequent removal of CSB and the stalled RNAPII from the damaged chromatin. While various mechanisms can potentially resolve RNAPII stalled at bulky lesions (Anindya et al., 2007; Bregman et al., 1996; Kleiman et al., 2005; Nakazawa et al., 2012; Ratner et al., 1998; Starita et al., 2005; Yasukawa et al., 2008), TC-NER emerges as the dominant pathway for its removal in p300-deficient cells.

**Figure 2.**
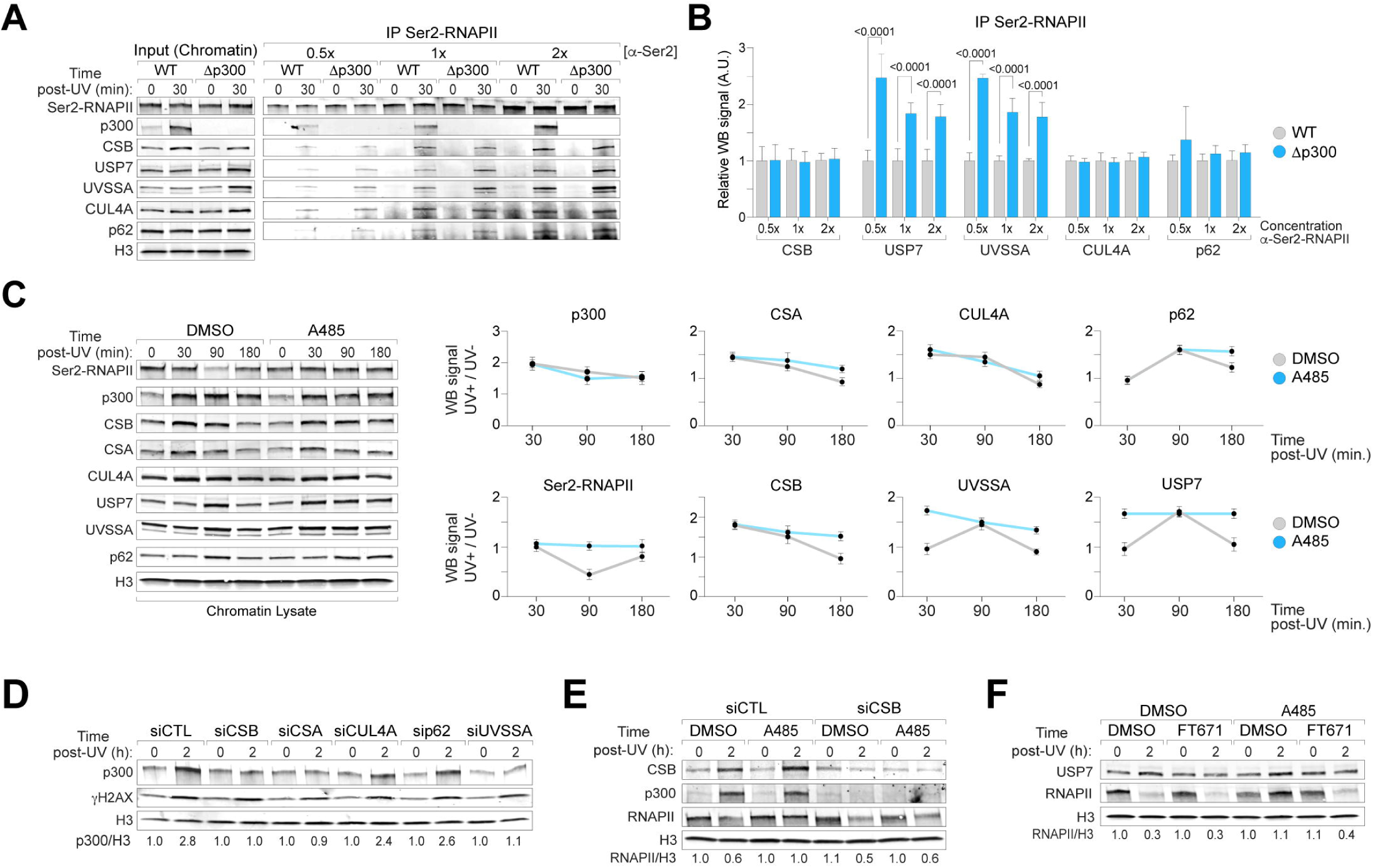
P300 controls UVSSA and USP7 interaction with RNAPII post DNA damage. **(A)** Western blot (WB) analyses for the indicated proteins of immunoprecipitations (IP) assays with different concentration of Ser2-RPB1 antibody (1X = 5 μg) at 0 and 30 minutes after UV-irradiation (20 J/m^2^) using chromatin-enriched extracts of WT or Δp300 HeLa cells. The input shown constitutes 3% of the chromatin fraction. **(B)** Quantification of the WB signals in *A*. P values from one-way ANOVA are provided. **(C)***Left*, WB analysis using the indicated antibodies and chromatin-enriched extracts of HeLa cells at different time points after UV irradiation (20 J/m^2^). DMSO or 1 mM A-485 was added to the culture 1 h before UV-damage. *Right*, relative quantification of the WB signals and their replicates. The relative values were calculated using H3 and normalized to the samples before UV damage across three independent replicates**. (D)** WB analysis of HeLa cells transfected with different siRNAs targeting different TC-NER factors and related genes. siRNA transfections were performed 24 h before UV irradiation (20 J/m^2^). The ratios of RNAPII and H3 signals relative to the samples before UV damage are shown. **(E)** WB analysis of HeLa cells transfected with siRNAs targeting CSB. siRNA transfections were performed 24 h before UV-irradiation (20 J/m^2^). Cells were incubated with DMSO or 1 μM A485 1 h before UV exposure (20 J/m^2^). The ratios of RNAPII and H3 signals relative to samples before UV damage are shown. **(F)** WB analysis from chromatin-enriched HeLa extracts at 0 and 2 h post-UV (20 J/m^2^). DMSO, 1 μM A-485 or 1 μM FT671 were added 1 h prior to UV exposure. The ratios of RNAPII and H3 signals relative to samples before UV damage are shown. All experiments were performed in at least three independent experimental replicates.

### P300 inhibits persistent transcription activity after DNA damage

Following exposure to UV irradiation, the stalling of the elongating RNAPII in gene bodies due to bulky DNA lesions results in a marked decrease of transcription at the 3’ end of the genes (Lavigne et al., 2017; Nakazawa et al., 2020; Tufegdzic Vidakovic et al., 2020). To assess the impact of p300 inhibition on RNAPII genome-wide distribution and activity after DNA damage, ChIP-seq analyses were conducted in both WT and p300-deficient cells, using antibodies against total RNAPII or Ser2-RNAPII. In the absence of DNA damage, RNAPII displayed the expected distribution pattern, featuring peaks near TSS, dispersion throughout gene bodies, and significant enrichment following transcription termination sites (TTS) (**Figures 3A, 3B, and S3A**). Also, Ser2-RNAPII exhibited its characteristic enrichment at the 3’ end of genes (**Figures 3C, 3D, and S3B**). Noteworthy, ChIP-seq normalized reads exhibited a strong correlation between independent experimental replicates throughout all experiments performed (**Figures S3C and S3D**). As previously observed (Lavigne et al., 2017; Nakazawa et al., 2020; Tufegdzic Vidakovic et al., 2020), two hours after UV exposure, both total RNAPII and Ser2-RNAPII distributions significantly decreased at the 3’ end of genes due to DNA damage-induced transcription stalling (**Figures 3A, 3B, 3C, 3D, S3A and S3B**). Total RNAPII also showed a decrease at TSSs and enhancers in WT cells following DNA damage (**Figures 3A, 3B, S3A and S3E**). These changes in RNAPII distribution correlated with reduced levels of RNAPII at chromatin and diminished 5-EU incorporation following DNA damage (**Figure 1**). Importantly, the impact of DNA damage on RNAPII was more pronounced in long genes compared to very short ones (0 - 5 kbp) (**Figure 3E and S3F**), as RNAPII traversing long genes had a higher probability of encountering bulky DNA lesions. Prior to DNA damage, p300-deficient cells exhibited similar profiles of total RNAPII and Ser2-RNAPII as WT cells. However, following UV irradiation, p300-deficient cells displayed a slight increase in total RNAPII and Ser2-RNAPII at the 3’ end of gene bodies and showed no changes at enhancers (**Figures 3A, 3B, 3C, 3D, S3A, S3B and S3E**). The observed high levels of total and Ser2-RNAPII at the 3’ end of genes suggested that transcription activity was not arrested in the gene body of genes in p300-deficient cells following DNA damage.

**Figure 3.**
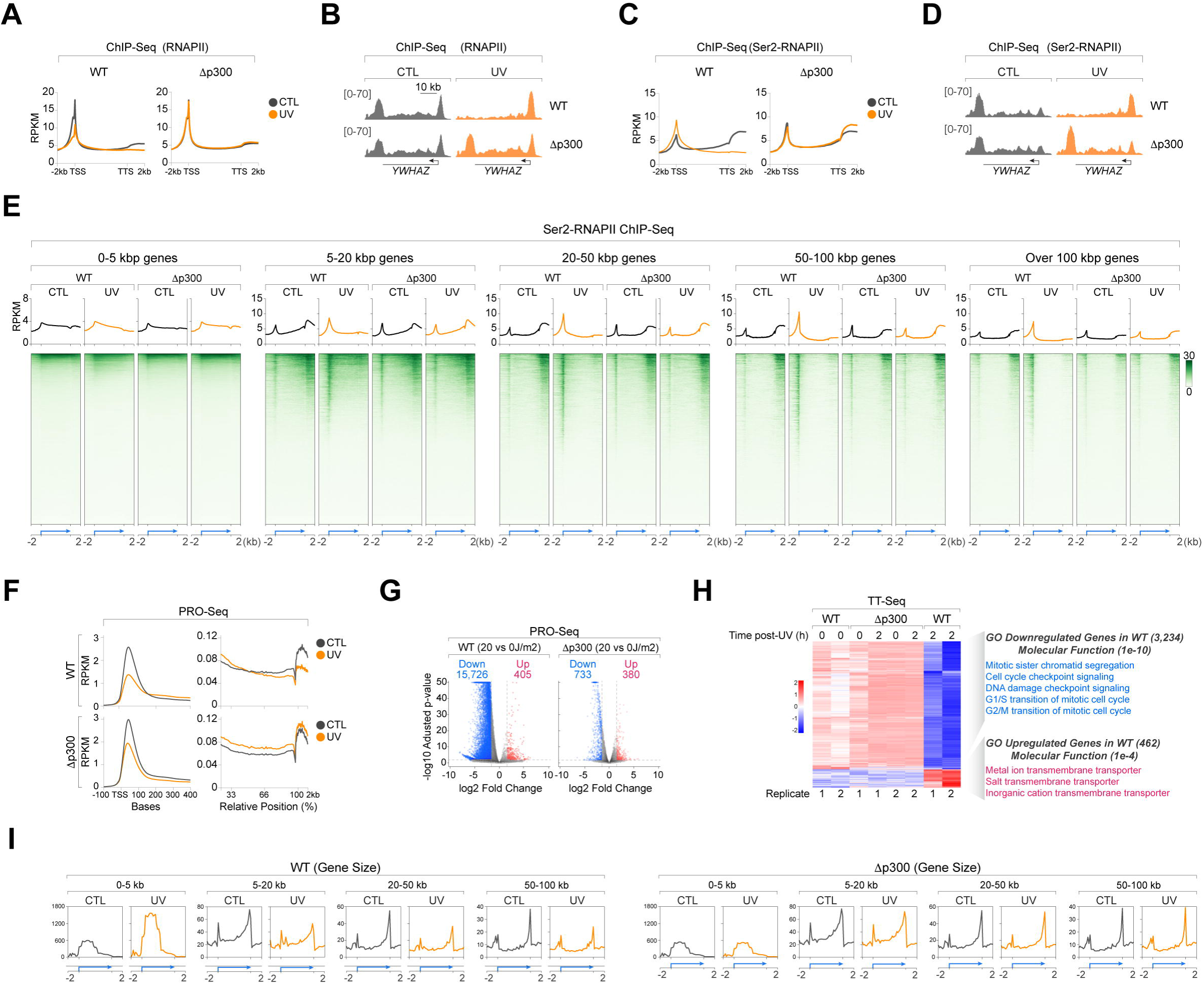
P300 inhibits persistent transcription activity after DNA damage. **(A)** Metagene analysis showing total RNAPII and Ser2-RNAPII occupancy measured by ChIP-seq at 0 and 2 h after UV-irradiation (20 J/m^2^) in either WT or Δp300 HeLa cells (RPKM, reads per kilobase of transcript per million mapped reads). **(B)** Representative genome browser examples of total RNAPII ChIP-seq. The x-axis indicates the chromosome position, and the y-axis represents normalized read density in reads per million [rpm]. **(C)** As in *A*, but for and Ser2-RNAPII. **(D)** As in *B*, but for Ser2-RNAPII. **(E)** ChIP-seq heatmap of Ser2-RNAPII shown in *C*, for different gene size clusters (0-5; 5-20; 20-50; 50-100; over-100 kilobase pair). **(F)** Metagene analysis showing distribution of PRO-seq signal at 0 and 2 h after UV-irradiation (20 J/m^2^) in both WT and Δp300 HeLa cells. **(G)** Volcano plots of genes with differential polymerase activity (log_2_FC ≥ 1.5, q-value < 0.05) derived from PRO-seq for 20 vs 0 J/m^2^. Only reads in the last 40% of the gene up to the transcription termination site (TTS) were used for the analysis. Samples were normalized according to their spike-in geometric mean. The number of downregulated genes is shown in blue, and the number of upregulated genes is shown in red. **(H)** *Left*, heatmap of log_2_ fold changes of nascent RNA-seq with 5-EU added 10 minutes before collection of samples from WT or Δp300 HeLa cells exposed to UV (20 J/m^2^). *Right*, gene-ontology (GO) enrichment analysis for downregulated and upregulated genes in WT cells exposed to UV from nascent RNA-seq data. The number of genes in each cluster is shown in brackets. **(I)** Metagene analysis showing total nascent RNA-seq data shown in *H*, for different gene size clusters. All experiments were performed in at least two independent experimental replicates.

To further explore this phenomenon, Precision Run-On Sequencing (PRO-Seq) was conducted in both WT and p300-deficient cells. PRO-Seq maps the location of actively engaged, non-arrested, or paused RNA polymerases with base-pair resolution (Kwak et al., 2013). In WT cells, RNAPII exhibited decreased transcription activity after UV damage, accumulating near TSS regions and diminishing at the 3’ end of genes due to RNAPII stalling (Lavigne et al., 2017; Nakazawa et al., 2020; Tufegdzic Vidakovic et al., 2020). Conversely, in p300-deficient cells, RNAPII displayed reduced transcription activity near the TSS region and an increase within gene bodies and 3’ end of genes after DNA damage (**Figures 3F and S3G**). Since in p300-deficient cells the protein levels of RNAPII do not change after DNA damage (**Figure 1**), these results suggested a widespread defect in promoter-proximal pausing RNAPII control in p300-deficient cells after UV irradiation. To confirm this observation, we determined the RNAPII traveling ratio (RTR) by measuring the occupancy of active RNAPII within the gene body in comparison to its promoter. For these analyses, the promoter was defined as 30 bp upstream and 300 bp downstream of the TSS, while the gene body was defined as the region from 300 bp downstream of the TSS to the first 20% of the gene body (**Figure S3H**). These regions were selected to measure the release of RNAPII before and after UV damage, ensuring that potential defects in transcription processivity in gene bodies and/or in transcription termination do not affect these analyses. These analyses suggested that after DNA damage, paused RNAPII is more readily released into gene-bodies in p300-deficient than WT cells.

To further characterize the impact of p300 deficiency on transcription control after DNA damage, we determined the number of genes affected in WT and p300-deficient cells. For this, we analyzed the PRO-Seq signal in the last 40% of the gene body, which differentiates genes where RNAPII stalled from those where it did not. Two hours after UV irradiation, WT cells showed a reduction in the transcription activity of 15,726 genes and an increase in 405 genes (**Figure 3G**). Whereas p300-deficient cells showed a reduction in only 733 genes and upregulation in 380 (**Figure 3G**). These findings were corroborated through transcription transcriptome sequencing (TT-seq) which provides a more accurate reflection of ongoing transcription than steady-state RNA-seq. In these analyses, WT cells exhibited a clear reduction in gene transcripts (3,234 transcripts) following UV irradiation (**Figure 3H**). Downregulated processes were predominantly linked to cell cycle checkpoints and DNA damage signaling, consistent with a stress response after UV damage (**Figure 3H**). Moreover, while WT cells exhibited a decrease in the expression of genes longer than 5 kb, they also showed upregulation of 462 genes shorter than 5 kb following UV damage (**Figures 3H and 3I**). This pattern has been previously observed, where short genes critical for survival after UV exposure are preferentially upregulated (Tufegdzic Vidakovic et al., 2020). Upregulated genes were mainly linked to transmembrane transporters (**Figure 3H**). However, p300-deficient cells displayed no significant alterations in gene transcript levels (**Figures 3H**). Similar results were obtained when analyzing RNA-seq data from chromatin-enriched fractions (**Figure S5I**). These results establish that, unlike WT cells, p300-deficient cells maintain high and productive transcription activity following DNA damage, which is consistent with the unchanged levels of RNAPII and 5-EU incorporation observed in p300-deficient cells after DNA damage (**Figure 1**).

### RNAPII bypasses blocking DNA lesions in p300-deficient cells

Upon UV exposure, transcription elongation is hindered due to RNAPII stalling at transcription blocking lesions (TBL) such as CPDs and 6–4PPs (Mayne and Lehmann, 1982) (Lavigne et al., 2017; Williamson et al., 2017) (Gyenis et al., 2014; Rockx et al., 2000). While two hours post UV irradiation WT cells exhibit the expected decrease in transcription activity due to stalling of RNAPII, p300-deficient cells showed high transcription activity along all the gene body (**Figure 3**). This suggests that RNAPII in p300-deficient cells can transcribe despite the presence of TBLs. To investigate this, we performed strand-specific ChIP-seq data analyses (Nakazawa et al., 2020; van der Weegen et al., 2021) for total and elongating RNAPII to assess its interaction with TBLs in both WT and p300-deficient cells post UV exposure (**Figure 4A**). If TBLs are present in the immunoprecipitated precipitated chromatin, the damaged DNA strand will amplify less efficiently due to the stalling of the DNA polymerase during the PCR step of the library preparation, indicating RNAPII stalled at a TBL (Nakazawa et al., 2020; van der Weegen et al., 2021). Consequently, in the case of an elongating RNAPII stalled at a TBL, we would expect a lower signal in the template strand compared to the coding strand. Expectedly, two hours post UV, WT cells showed reduced signal for total and elongating RNAPII in the template strand but not in the coding strand (**Figures 4B-4E**). To assess the presence of TBLs in specific genes, we calculated the Strand-Specificity Index (SSI) (Nakazawa et al., 2020; van der Weegen et al., 2021) (**Figure 4F**). This index is determined by the ratio of read counts mapped in both the forward and reverse directions within the gene’s body. The SSI values, which may be either positive or negative, indicate not only the bidirectional orientation of genes but also the relative proportion of TBLs that persist on the transcribed DNA strand as compared to the non-transcribed strands. In these analyses, most transcribed genes in WT cells showed damage in the template strand two hours after UV damage (**Figure 4F**). However, p300-deficient cells exhibited no strand bias for total- or Ser2-RNAPII after UV, suggesting that RNAPII in these cells does not pause at TBLs (**Figures 4B-4F**). To corroborate these findings and to further assess the ability of RNAPII to transcribe in the presence of TBLs, we evaluated how RNAPII travels through UV-damaged chromatin in both WT and p300-deficient cells. To achieve this, we began by suppressing RNAPII transition from the release to the elongation phase by treating cells with 5,6-dichlorobenzimidazole 1-beta-D-ribofuranoside (DRB) for 3 hours. Following this treatment, the cells were split into two groups: one group was exposed to UV irradiation, while the other group (CTL) was not irradiated. Five minutes after the UV exposure, DRB was removed from both groups of cells to allow transcription re-start. Samples were collected after DRB arrest (0 minutes) and 45 minutes post UV irradiation for ChIP-seq analyses of total and Ser2-RNAPII. Following DRB exposure, total and Ser2-RNAPII levels decreased at chromatin fractions and accumulated at the promoter area, indicating transcription arrest in both WT and p300-deficient cells (**Figures 5A, and S4A-S4E**). Post DRB release, total and Ser2-RNAPII returned to normal levels at the chromatin with typical distribution in genes of various lengths in non-damaged WT and p300-deficient cells (**Figures 5A, and S4A-S4E**). However, in irradiated WT cells, total and Ser2-RNAPII predominantly accumulated at the promoter-proximal region, while p300-deficient cells exhibited high total and Ser2-RNAPII levels throughout gene bodies and 3’ ends, including in long genes (>100 kb) like *PHLDB2* and *TLE4* following UV damage (**Figures 5A, and S4A-S4E**). In addition, we combined 5-EU metabolic labeling of newly synthesized RNA with DRB chase experiments to measure RNAPII activity through damaged chromatin. These studies further demonstrated that, unlike in WT cells, RNAPII in p300-deficient cells exhibits high activity levels through damaged chromatin (**Figure 5B**). These results suggest that RNAPII transcribes through damaged chromatin either by bypassing the blocking lesions or, after stalling, by backtracking to allow repair of the TBL before continuing with elongation. To distinguish between these two possibilities, we performed strand-specific ChIP-seq analyses before and after UV irradiation in cells exposed to DMSO or A-485. During the first two hours post UV, control cells showed an increased strand bias and reduced RNAPII protein levels and occupancy (**Figures 5C and S5A-S5C**). These effects induced by UV lessened after three hours, indicating DNA repair activity (**Figures 5C, S5A and S5B**). In contrast, A-485 exposed cells, even 3 hours after UV, showed minimal changes on RNAPII occupancy, strand bias, and protein levels (**Figures 5C, S5A and S5B**). Yet, when A-485 was removed one hour after UV, the response of RNAPII mirrored that observed in damaged control cells, showing that p300 inhibition removal reinstates RNAPII stalling at TBLs. Indeed, after removal of A-485, cells showed a decreased RNAPII occupancy and protein levels, and increased strand bias (**Figures 5C, S5A and S5B**). These data demonstrated that RNAPII transcribes through damaged chromatin by bypassing TBLs instead of backtracking to allow TBL repair.

**Figure 4.**
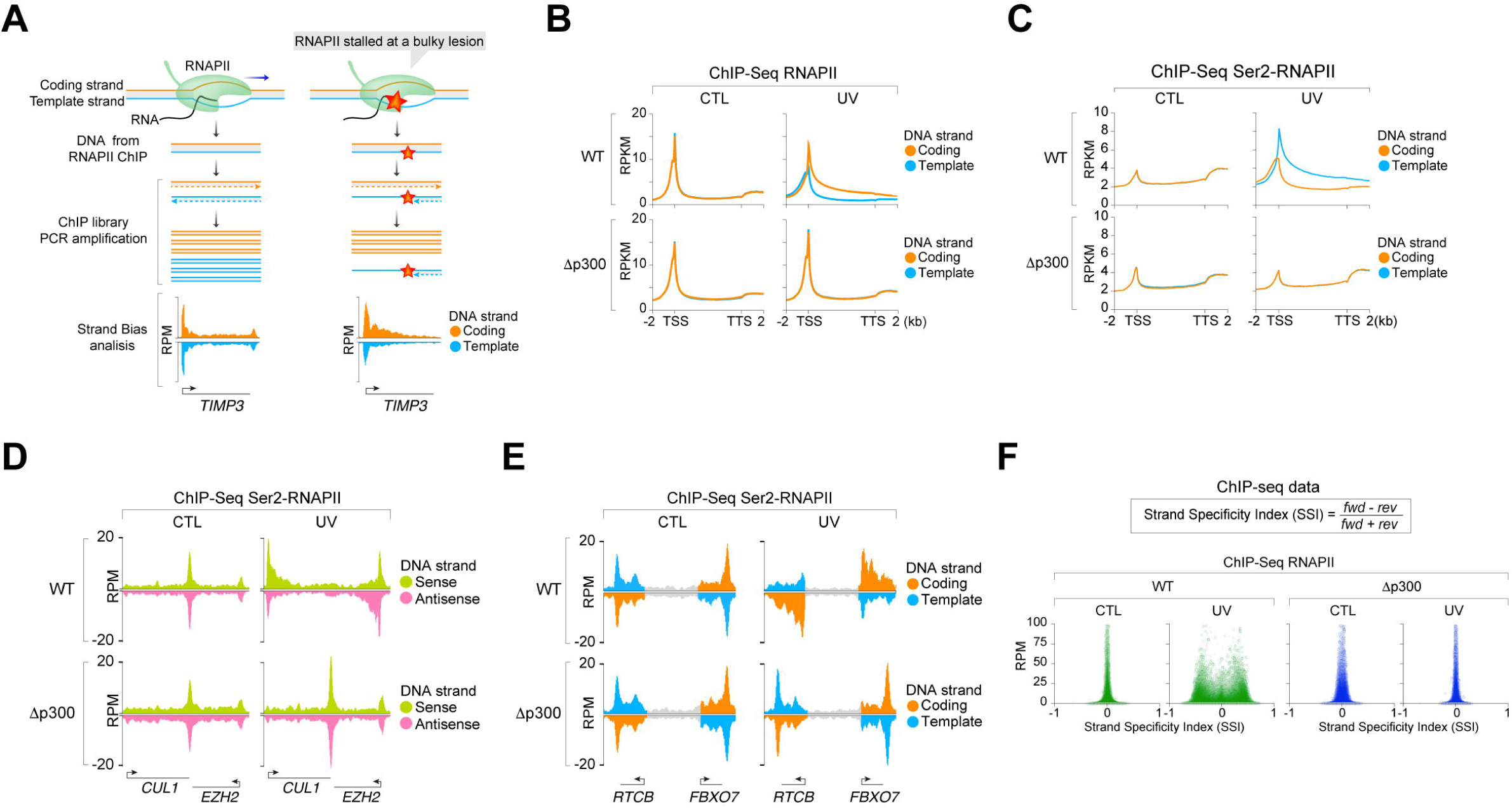
P300 inhibits the bypass of blocking DNA lesions by RNAPII. **(A)** Schematic representation of the strand-bias ChIP analysis. **(B)** and **(C)** Metagene analysis showing total RNAPII or Ser2-RNAPII occupancy on the coding or template DNA strand, measured by ChIP-seq at 0 and 2 h after UV-irradiation (20 J/m^2^) in either WT or Δp300 HeLa cells. **(D)** Sense (green) and antisense (pink) strand-biased ChIP-seq signals of the same data as in *B* are shown for representative convergent genes**. (E)** Coding (orange) or template (light blue) strand-biased ChIP-seq signals of the same data as in *C* are shown for genes oriented in opposite directions. **(F)** The SSI was defined to evaluate strand bias in RPB1-ChIP DNA fragments and calculated based on the number of forward and reverse reads within each gene body. Positive and negative values reflect gene orientation. A unimodal SSI distribution is seen in control samples, whereas a bimodal pattern emerges 2 h after UV exposure. SSI was computed using normalized signal from the coding and template strands of ChIP-seq data, before (CTL) or 2 h after UV irradiation (20 J/m²), in WT and Δp300 HeLa cells. Genes with <1 RPM were excluded. All experiments were performed in at least two independent experimental replicates.

**Figure 5.**
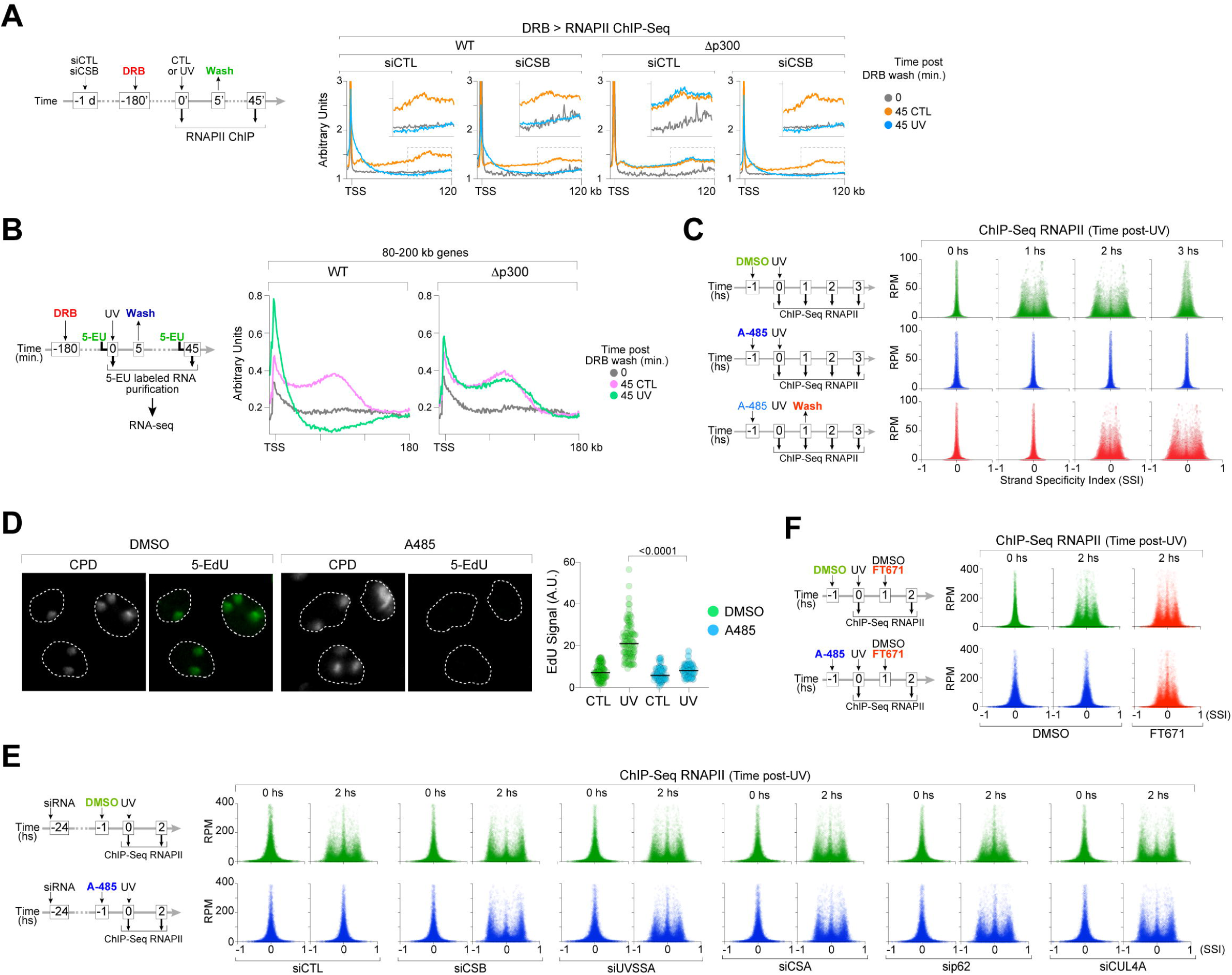
TC-NER and USP7 facilitate the bypass of blocking DNA lesions by RNAPII in p300-deficient cells. **(A)** *Left,* schematic of the experimental strategy. *Right,* Metagene analysis showing RNAPII distribution from ChIP-seq data derived from the DRB chase experiment. WT and Δp300 HeLa cells were transfected with siRNA control (siCTL) or against CSB (siCSB) 24 h before exposure to UV (20 J/m^2^). Samples were filtered using IgG control, and only genes with a size between 80 and 200 kb were used for the analysis. **(B)** *Left,* schematic of the experimental strategy. *Right,* metagene analysis showing 5-EU labeled RNA signal in WT or Δp300 HeLa cells exposure to UV (20 J/m^2^). 5-EU (1 mM) was added 10 minutes before sample collection. Samples were filtered using ‘No 5-EU’ control, and only genes with a size between 80 and 200 kb were used for the analysis. **(C)** *Left*, schematics showing the experimental strategy. *Right*, scatter plots of strand-specific index (SSI) for each gene in the indicated conditions post-UV irradiation (20 J/m^2^). **(D)** *Left*, representative images TC-NER-specific unscheduled DNA synthesis (UDS) in HeLa cells transfected with siRNA against XPC. Scale bar, 10 μm. Cells were irradiated with 100 J/m^2^ UV through 5 μm pore membranes and exposed to DMSO or A-485 (1 μM) one hour before UV. *Right*, quantification of the 5-EdU levels in the damaged areas of the cells 3 hours post UV exposure. Each datapoint represents the UDS signal from a single cell from three independent replicates. The bar represents the mean of all data points. Scale bar, 10 μm. *p*-values by one-way ANOVA tests. **(E)** *Left*, schematics of the experimental strategy. *Right*, scatter plots of SSI for each gene in the indicated conditions post-UV irradiation (20 J/m^2^). **(F)** *Left*, schematics of the experimental strategy. DMSO, 1 μM A-485 or 1 μM FT-671 were added 1 h prior to UV exposure. *Right*, scatter plots of SSI for each gene in the indicated conditions post-UV irradiation (20 J/m^2^). All experiments were performed in at least two independent experimental replicates.

We next conducted base-resolution mapping of stalled RNAPII using ChIP-seq data, both before and after UV irradiation. This analysis aimed to ascertain whether RNAPII accumulates near CPDs post UV exposure (Nakazawa et al., 2020). Our findings indicate an expected rise in the number of mapped reads on the coding strands proximal to GpA, ApA, GpG, and ApG dinucleotides, which was not observed for TpT, CpC, ApT dinucleotides in control (DMSO) cells after UV exposure (**Figure S5D**). This increase is attributed to the stalling of RNAPII at CPDs produced by UV irradiation, on the template strand (C<>T, T<>T, C<>C, and T<>C respectively). In control cells this enrichment was evident in the initial two hours, then diminished an hour later, likely due to DNA repair (**Figure S5D**). Importantly, this accumulation near CPDs was absent in cells continuously exposed to A-485. However, in cells where A-485 was removed an hour post UV irradiation, we observed an increase in mapped reads preferentially near CPDs in the coding strand at 2 hours post UV, which increased over time (**Figure S5D**). These observations suggest that upon the removal of A-485, RNAPII stalls at TBLs in a way similar to control cells. These results also suggest that TC-NER does not repair blocking lesions in p300-deficient cells. Supporting this, a TC-NER-mediated repair synthesis assay showed that while WT cells exhibit DNA synthesis two hours after UV damage, p300-deficient cells display impaired repair synthesis (**Figure 5D**).

### Bypass of blocking DNA lesions by RNAPII in p300-deficient cells depends on TC-NER and USP7

Next, we sought to determine whether TC-NER plays a role in the bypass of TBLs by RNAPII in p300-deficient cells. To test this, we inhibited several TC-NER factors in WT and p300-deficient cells using siRNAs (**Figure S6A**). Although inhibition of CSB, CSA, UVSSA, and p62 did not affect RNAPII genome-wide distribution or strand bias before or after UV damage in WT cells, their inhibition blocked the bypass of TBLs by RNAPII in p300-deficient cells after UV damage (**Figures 5E and S6B**). Moreover, inhibition of USP7 with FT-671 one hour after UV damage increased strand bias and prevented RNAPII from reaching the 3’ end of genes in p300-deficient cells (**Figures 5F and S6C**). Additionally, USP7 inhibition rescued the transcription shutdown defect observed in p300-deficient cells, highlighting the opposing roles of p300 and USP7 in regulating transcriptional activity after DNA damage (**Figure S6D**). Likewise, inhibition of CSB blocked RNAPII elongation over damaged chromatin in p300-deficient cells (**Figures 5A, S4D and S4E**). Altogether, these findings show that the bypass of TBLs by RNAPII in p300-deficient cells depends on TC-NER and USP7.

### P300-deficient cells produce functional mRNAs after DNA damage

To investigate whether the persistent transcription activity observed in p300-deficient cells after DNA damage produces functional mRNAs, we combined 5-EU metabolic labeling of newly synthesized RNA with RNA-seq from polysome-enriched fractions. First, we sequenced RNA samples from WT and p300-deficient cells before and after UV damage. In these analyses, WT cells exhibited a significant decrease in 1,551 transcripts and an increase in 118, whereas p300-deficient cells showed a decrease in only 371 transcripts and an increase in 668 (**Figures S7A and S7B**).

Next, we evaluated the distribution of 5-EU-labeled transcripts in polysome-enriched fractions of WT and p300-deficient cells before and after UV damage. Consistent with the global transcriptional shutdown observed in WT cells, 5-EU-labeled transcripts decreased markedly two hours post-UV damage (**Figure S7C**). In contrast, p300-deficient cells exhibited no significant differences under the same conditions (**Figure S7C**). Notably, no significant differences in splicing or mutation patterns were observed in chromatin-enriched or polysome-enriched mRNAs from WT and p300-deficient cells following DNA damage (**Figures S8A-S8E**).

These findings demonstrate that in p300-deficient cells, RNAPII remains engaged in productive transcription even in the presence of unrepaired TBLs. This leads to the complete transcription of genes into functional mRNAs, highlighting the ability of RNAPII to bypass these lesions and maintain effective full-length transcription.

### P300 inhibition increases the toxicity of UV radiation in a USP7-dependent manner

Next, we examined the impact of p300 inhibition on cellular proliferation following exposure to UV irradiation. Colony assays revealed a high sensitivity of Δp300 HeLa cells to UV irradiation when compared to WT HeLa cells (**Figure 6A**). Similarly, A-485 demonstrated remarkable synergy with UV irradiation in killing U2OS cells (**Figure 6B**). Importantly, this synergistic effect was achieved with just 8 hours of exposure to A-485. This reduction in cell proliferation capacity coincided with an increase in DNA damage and apoptosis (**Figures 6C and S9**). These data highlighted the importance of p300 activity for survival in the first hours post DNA damage with UV radiation.

**Figure 6.**
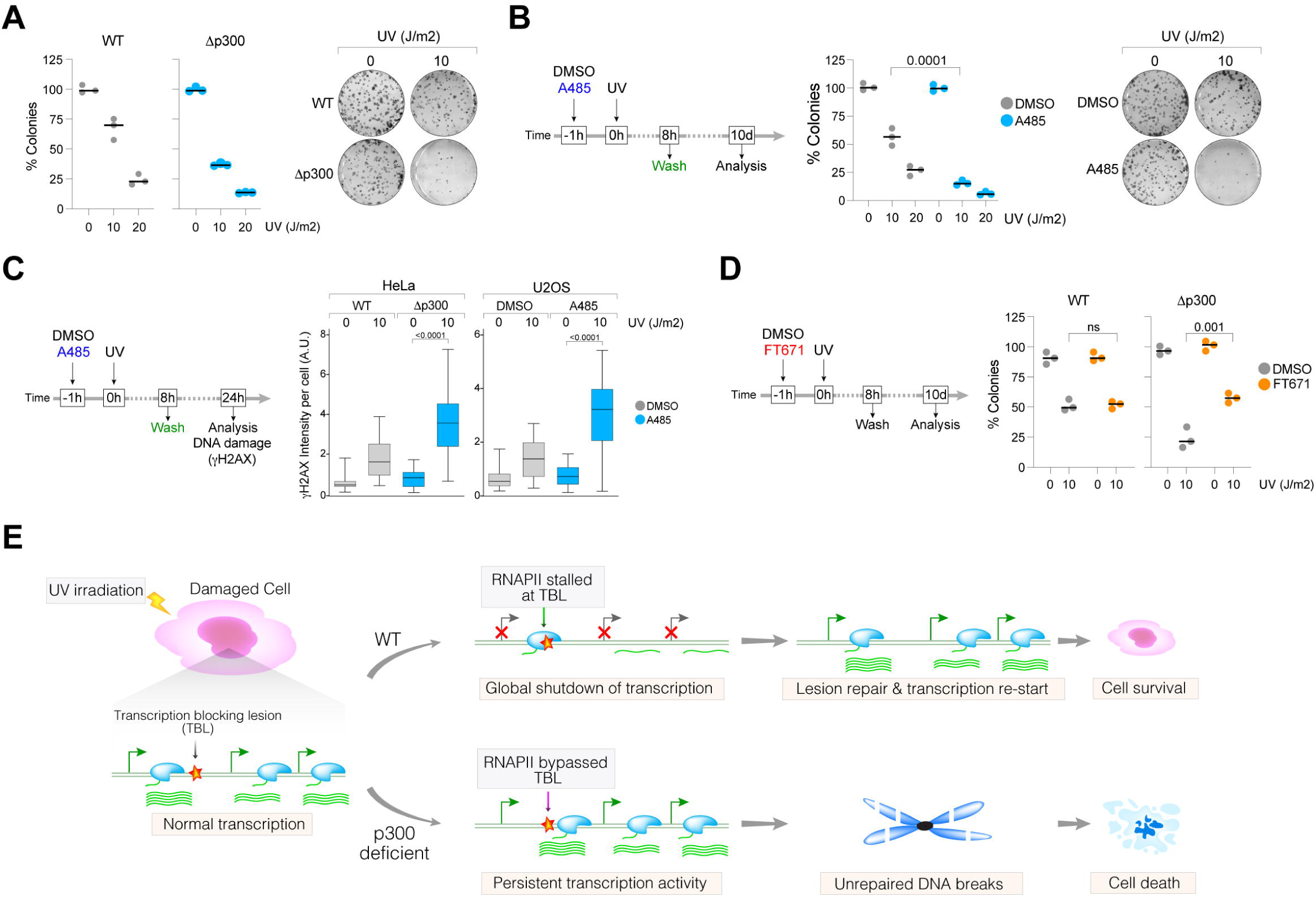
Inhibition of p300 enhances UV-induced toxicity through a mechanism that depends on USP7. **(A)** *Left*, percentage of colonies in WT and Δp300 HeLa cells before or after exposure to different doses of UV-irradiation (0-10-20 J/m^2^). *Right*, representative images of colony formation assays from WT and p300-deficient HeLa exposed to UV radiation. **(B)** As in *A*, but using U2OS cells. DMSO or 1 μM A-485 were added to the culture 1 h before UV-damage and removed 8 h after UV exposure. **(C)** *Left*, schematic of the experimental design. *Right*, quantification of γH2AX signal intensity per cell in the indicated cell lines 0 and 24 h after exposure to UV radiation (10 J/m2). A-485 (1 μM) or DMSO were added to the culture together one hour before UV radiation and both were removed 8 h post UV. *p*-values by one-way ANOVA tests. One hundred cells were analyzed per condition/experiment. **(D)** *Left*, schematic of the experimental design. *Right*, quantification of colony assays with WT and Δp300 HeLa cells in presence of DMSO or 0.25 μM USP7 inhibitor (FT671). All compounds were removed 8 h later as indicated in the schematic. All experiments were performed in at least three independent experimental replicates. **(E)** Proposed model. Simplified schematic illustrating the consequences of RNAPII overcoming transcription blocking lesions (TBLs) generated by UV irradiation and maintaining elevated levels of transcription following DNA damage. Our findings reveal that blocking p300 allows RNAPII to circumvent TBLs, resulting in continuous transcription activity and increased DNA damage that affects cell proliferation.

Next, while inhibition of USP7 with FT671 had no effect on cell proliferation or the toxicity of UV alone, it nearly completely mitigated the increased toxicity caused by the combination of p300-deficiency with UV radiation (**Figure 6D**). These results align with the rescue of defective RNAPII degradation and stalling at blocking lesions observed in p300-deficient cells following TC-NER or USP7 inhibition (**Figures 2F, 5F, S6C and S6D**). These results underscore the significance of the p300-UVSSA-USP7 axis in controlling RNAPII degradation following DNA damage and its crucial role in cell survival. Moreover, these data demonstrate that the toxicity of p300 inhibition when combined with UV is TC-NER dependent.

In summary, these findings provide strong evidence that inhibiting the bypass of transcription-blocking lesions by RNAPII is crucial for cell survival following DNA damage.

## Discussion

The implications of a genome-wide transcription shutdown after high DNA damage levels have been thoroughly investigated (Doetsch, 2002; Lans et al., 2019; Noe Gonzalez et al., 2021). However, it is not yet clear how cell survival is impacted when cells evade this DNA damage-induced transcription response. In this report, we explore the consequences of RNAPII bypassing TBLs and sustaining high transcription activity after DNA damage. We demonstrate that inhibiting p300 allows RNAPII to bypass TBLs, leading to persistent transcription activity and DNA damage that compromises cell survival (**Figure 6E**). This bypass of TBLs by RNAPII reveals a novel cellular stress post-DNA damage.

We show that p300 plays a crucial role in removing the stalled RNAPII from damaged chromatin. Our studies suggest that this is most probably due to p300 inhibiting the interaction between USP7 and the TC-NER complex. Consequently, p300 inhibition enhances the interaction of USP7 and UVSSA with TC-NER and paused RNAPII upon encountering a blocking lesion, preventing the removal of CSB and RNAPII from damaged chromatin and thereby facilitating the bypass of TBLs. Supporting this, the inhibition of CSB or USP7 rescues RNAPII degradation, the stalling of RNAPII at TBLs, and the proliferation defects caused by p300 deficiency in cells exposed to DNA damage. This also underscores that, despite the involvement of p300 in different cellular processes -such as proliferation, differentiation, apoptosis, and DNA repair (Bannister and Kouzarides, 1996; Dancy and Cole, 2015; Ogryzko et al., 1996)-during the first hours following DNA damage induced by UV irradiation, its primary role is mediated through the TC-NER pathway.

While p300 inhibition prevents the ubiquitylation and degradation of RNAPII after DNA damage, this alone does not appear to be sufficient to facilitate the bypass of TBLs by RNAPII. Notably, we show that while inhibition of CUL4A blocks RNAPII removal from the damaged chromatin, it does not allow the bypass of TBLs. Moreover, a mutant form of RPB1 (RPB1K1268R), which is resistant to ubiquitylation by CRL4^CSA^ and degradation after DNA damage, still exhibits a halt in transcription elongation (Nakazawa et al., 2020; Tufegdzic Vidakovic et al., 2020). Interestingly, following DNA damage, CSB and CSA interact with stalled RPB1K1268R, but the absence of TFIIH interaction shows that only partial recruitment of the TC-NER complex occurs (Nakazawa et al., 2020). This observation aligns with the finding that the binding of UVSSA to the lesion site, which is dependent on the ubiquitylation of K1268, ultimately facilitates the recruitment of TFIIH (Kokic et al., 2024; Nakazawa et al., 2020; Okuda et al., 2017; van der Weegen et al., 2020). This contrasts with the scenario in p300-deficient cells, where, in addition to CSB, CSA, UVSSA, and TFIIH interacts with the elongating RNAPII after DNA damage. However, despite this TC-NER recruitment, TBLs remain unrepaired in p300-deficient cells. This observation raises some interesting points. Firstly, even though in p300-deficient cells RNAPII continues transcribing in the presence of TBLs, it likely pauses at these lesions as TC-NER proteins associate with elongating RNAPII. This TC-NER/RNAPII interaction in p300-deficient cells can be observed even at timepoints post DNA damage where WT cells decrease TC-NER recruitment to the damaged chromatin. This is most probably due to the presence of unrepaired TBLs in p300-deficient cells at later timepoints as shown here. Secondly, although p300-deficient cells show a lower level of ubiquitylation of RNAPII after DNA damage, some ubiquitylation of K1268 needs to take place to allow UVSSA and TFIIH recruitment (Fei and Chen, 2012; Groisman et al., 2006; Higa et al., 2018; Kokic et al., 2024; Nakazawa et al., 2020; Nakazawa et al., 2012; Schwertman et al., 2012; van der Weegen et al., 2021; van der Weegen et al., 2020; Zhang et al., 2012). Third, TC-NER not only contributes to RNAPII removal from damaged chromatin but also plays a role in the bypass of TBLs. Fourth, although RNAPII can bypass TBLs -potentially allowing lesion access and repair by the recruited TC-NER-this does not appear to occur efficiently. It will be important to determine whether other factors involved in TBL bypass interfere with TC-NER–mediated repair. While further research is needed to clarify the mechanisms by which RNAPII bypasses these lesions and to identify the p300 acetylation targets involved, our findings highlight TC-NER and USP7 as central components of this process.

We also demonstrate that degradation of the stalled RNAPII is less toxic for the cell than bypassing the TBLs and continuing transcription. This indicates that cells prefer to degrade the elongating RNAPII and disassemble the transcription machinery, followed by a global transcription shutdown, rather than risk the chance of TBL bypass and a persistent transcription activity. This is most likely due to the untimely production of proteins after DNA damage in cells that bypass TBLs, which could compromise cell survival.

Finally, although our work primarily focuses on dividing cells, it may also have implications for non-dividing cells. Most defects observed in human cells with a defective TC-NER pathway are associated with an impaired recovery of transcription activity due to defective TC-NER recruitment or a depleted RNAPII pool after DNA damage (Kokic et al., 2024; Tufegdzic Vidakovic et al., 2020). Our studies reveal that inhibiting p300 hinders TC-NER activity without depleting the RNAPII pool after DNA damage, which hints at the possibility of previously unrecognized human syndromes associated with TC-NER deficiencies.

## Acknowledgments

We are indebted to members of the Verdun and Shiekhattar laboratory for discussions.

## Author Contributions

R.E.V., C.P.B., and L.D.C. designed the study and analyzed the experiments. C.P.B. conducted most of the experiments. L.D.C. conducted most of the RNA-seq and ChIP-seq analyses with C.P.B. conducting part of the analyses. P.R.C. performed the PRO-seq experiments and F.P. the PRO-Seq bioinformatic analyses in the laboratory of R.S.. R.E.V. and C.P.B. supervised the experiments and provided intellectual support toward the interpretation of results. R.E.V. wrote the manuscript. All authors commented on the manuscript.

## Data and materials availability

All data needed to evaluate the conclusions in the paper are present in the paper and/or the Supplementary Materials. All raw and processed NSG data were deposited in the NCBI Gene Expression Omnibus under the GEO accession numbers GSE261052 (for ChIP-seq data), GSE261053 (for PRO-seq data), and GSE261054 (for RNA-seq data).

## Materials and Methods

### Cell lines and culture conditions

The following cell lines were used in this study: HeLa, human cervical cancer cell line (ATCC, #CCL-2); U2OS, human bone osteosarcoma epithelial cell line (ATCC, #HTB-96); IMR90, normal human lung fibroblast cell line (ATCC, #CCL-186). HeLa and U2OS cells were maintained in DMEM (Gibco, #11965-092) supplemented with 10% fetal bovine serum (FBS, Hyclone, #SH30910), 1X GlutaMAX (Gibco, #35050-061), 1X MEM non-essential amino acids (Corning 25-025-Cl) and 1X Antibiotic Antimycotic solution (Corning, #30-004-Cl), 5% CO_2_, and 21% O_2_. IMR90 cells were maintained in the similar growth conditions but with 15% FBS, and 3% O_2_. To generate Δp300 and ΔUSP7 HeLa cell lines, we employed CRISPR/Cas9-based gene editing strategy. For deletion of p300 we used the p300 CRISPR/Cas9 KO Plasmid (Santa Cruz, #SC400055) which produced a deletion from exon 4 to exon 29. For deletion of USP7 we used the HAUSP CRISPR/Cas9 KO Plasmid (h2) (Santa Cruz, #SC-402013-KO-2).

### Ultraviolet C (UVC) irradiation and chemical treatments

UVC-irradiation was performed in a UV-crosslinker (VWR, #89131-484) and the given doses (5, 10, or 20 J/m^2^) were monitored with a UV-meter (EXTECH, #SDL470). Cells were treated with A-485 (MCE, #HY-107455), FT-671 (MedKoo, #471006), Cycloheximide (Thermo Fisher Scientific, #J66901-03), MG132 (Sigma-Aldrich, #M8699), DRB (Sigma-Aldrich, #D1916), at the concentrations specified in each figure legend, DMSO (Sigma-Aldrich, #D8418) was used as vehicle control. All drugs were suspended in DMSO. For most experiments, drug administration was performed 1 h before UV-irradiation, unless otherwise noted.

### Transfection of siRNA

Cells were seeded into 100 mm plates at 1 x 10^6^ the day prior to siRNA transfection. Short-interfering RNAs (siRNAs) (Dharmacon: SMARTPool: #D-001810-10-20 for siCTL, #L-003486-00-0020 for siP300, #L-004888-00-0020 for siCSB, #L-012610-00-0020 for siCUL4A, #L-010924-00-0020 for siP62, #L-024139-02-0020 for siUVSSA, #L-011008-00-0020 for siCSA, #L-003477-00-0020 for CREBBP/CBP), and #L-016040-00-0005 for siXPC were transfected using jetPRIME transfection reagent (Polyplus) following manufacturer instructions.

### Cell proliferation and colony forming assay

For cell proliferation assays, 1x10^4^ cells were seeded into 6-well plates. Cell number was counted with a Countess II Automated Cell Counter (Invitrogen) on days 0, 2, 4, 6, and 8 for IMR90 cells or 0, 1, 2, 3 and 4 for U2OS and HeLa cells. For comparison of clonogenic capacity, 2000 cells were seeded into 100 mm plates. 10 days after plating, cells were washed twice with 1x PBS and fixed with cold 100% methanol at 4 °C for 20 min. Colonies were then stained with crystal violet solution (25% methanol with 0.5% w/v crystal violet, Sigma Aldrich # C6158-50G,) at room temperature for 30 min. After staining, the plates were gently washed with water and air-dried. Colonies were imaged with an Epson V750 Pro photo-scanner. Colonies with >50 cells were scored as surviving fraction. Synergy was estimated with *SynergyFinderPlus* software (https://synergyfinder.org). A combination index value >10 indicates synergism, values between 0 and 10 indicate additive and values < 0 indicate antagonism.

### Cellular fractionation lysates

The procedure begins with the washing of cells with 1x PBS, followed by resuspending the pellet in *Buffer 1* (10 mM Tris-HCl pH 8, 1.5 mM MgCl_2_, 10 mM KCl, 0.05% IGEPAL, 0.5 mM DTT and protease inhibitors) and incubating it on ice for 10 min. Subsequent centrifugation at 500 xg for 5 min at 4 °C separates the cytoplasmic fraction, which was stored at -20 °C. The pellet was then resuspended in Buffer 1 and centrifuged again to remove residual cytoplasmic components. The nuclear pellet was resuspended in *Buffer 2* (5 mM Tris-HCl pH 8, 1.5 mM MgCl_2_, 0.2 mM EDTA, 26% Glycerol, 0.5 mM DTT and protease inhibitors), followed by centrifugation at 10,000 xg for 20 min at 4 °C to collect the nucleoplasm fraction, stored at -20 °C. The remaining pellet, which corresponds to the chromatin-enriched fraction, was resuspended in *Buffer 3* (50 mM Tris-HCl pH 8, 300 mM NaCl, 10% glycerol, 0.2% IGEPAL, and protease inhibitors) and subjected to sonication cycles with a Bioruptor for 5 min in 30 seconds ON-OFF cycles. After a final centrifugation at maximum speed for 20 min, the supernatant was designated as the soluble chromatin fraction and stored at -20 °C. The remaining pellet was resuspended in RIPA buffer with protease inhibitors, subjected to additional sonication cycles, and labeled as the insoluble chromatin fraction. Quantification was performed via BIO-RAD DC Protein Assay.

### Immunoprecipitation (IP) assay

IP experiments were performed using chromatin-enriched extracts. Briefly, cells were lysed as described in the cellular fractionation section. For each IP assays, 500 μg of protein from the chromatin-enriched fraction was incubated overnight with 5 μg of antibody or IgG followed by 50 μl of Protein A/G PLUS agarose beads (Santa Cruz, #sc-2003) for 2 h. For quantitative IP experiments targeting Ser2-RNAPII, three antibody concentrations were evaluated: 2.5 µg, 5 µg, and 10 µg (corresponding to 0.25x, 0.5x, and 1x, respectively). DNAseI (20 μg/mg of protein), and Ethidium bromide (10 μg/ml) (Sigma-Aldrich, #E1510) were added to decrease DNA-dependent protein association. IP material was washed three times with high-salt buffer (50 mM Tris-HCl, 500 mM NaCl, 1% Triton X-100) and eluted with Laemmli buffer (BIO-RAD, #1610737EDU), then loaded for SDS-PAGE. For IP experiments with A-485 or USP7, cells were treated with DMSO or the respective inhibitor (1 μM) 1 h before UV exposure (20 J/m²).

### Western blotting

Western blotting was performed using standard protocols. Cell lysates were run on gradient (4-15%) SDS-PAGE gel, transferred onto a nitrocellulose blotting membrane (Cytiva, #10600002). Non-specific binding was blocked with 3% skim milk in Tris-buffered saline with 0.1% Tween-20 (TBS-T), followed by overnight incubation with primary antibodies. List of antibodies utilized in the study include those targeting RNA polymerase II and associated factors, histones, DNA damage response proteins, and various transcriptional regulators. For detection of RNA polymerase II, antibodies against Rpb1 NTD [D8L4Y] and CTD [4H8] (Cell Signaling Technology, #14958 and #2629), as well as phospho-specific antibodies recognizing Ser2 and Ser5 of the CTD repeat (Abcam, ab5095 and ab5131) were used. Anti-p300 antibodies from Active Motif (#61401), Abcam (ab14984), and Santa Cruz (sc-32244) were employed, along with anti-CBP from Thermo Fisher (PA5-27369). DNA damage was monitored using γ-H2AX antibodies from Millipore (clone JBW301, 05-636) and Cell Signaling Technology (#2577). Additional antibodies were against anti-ubiquityl-Histone H2A (Lys119) [D27C4] (CST, #8240), histone H3 (Abcam, ab4729), and acetylated histone variants H3 (K27) and H2B (K15) from Diagenode (C15310139 and C15410220); Histone H3 [1G1] (Santa Cruz, sc-517576); gamma-tubulin antibodies from GeneTex (GTX11316) and Sigma-Aldrich (T5192); anti-Vinculin [EPR20407] (Abcam, ab219649), anti-CSA [EPR9237] (Abcam, ab137033), anti-CSB [EPR9237] (BETHYL, A301-345A), and anti-UVSSA [GT816] from Thermo Fisher (MA5-27844) and GeneTex (GTX629742); GTF2H1 (p62) was detected with Abcam (ab232982), and USP7 and Cullin 4A (Cul4A) with Thermo Fisher antibodies (PA5-34911 and PA5-17101, respectively). Normal Rabbit IgG from Cell Signaling Technology (#2729) was used as a control. Secondary antibodies: IRDye 800CW donkey anti-mouse IgG (LI-COR, #926-32212) and IRDye 800CW donkey anti-rabbit IgG (LI-COR, #926-32213). WB signals were imaged on an Odyssey CLx imaging system (LI-COR).

### MultiDSK purification

The MultiDSK plasmid was developed from the yeast Dsk2 ubiquitin-binding domain (amino acids 327-373). This sequence was duplicated five times in a row, with a 7 amino acid linker (GAGSAGA) included between each duplicate. The sequence was synthesized (Azenta Life Sciences), and inserted into the GST fusion expression vector pGEX-4T1, resulting in the creation of pGST-MultiDSK. BL21(DE3) competent bacteria were transformed with the GST-MultiDSK plasmid and a single colony selected. The GST-MultiDSK fusion protein expression was initiated by inducing a culture (0.6 OD, 200 ml in terrific broth media) with IPTG (1 mM), followed by an incubation at 30 °C for 4 h. For optimal results, bacteria were either pelleted and frozen, with a quick room temperature defrost the next day, or processed on the same day. Twenty pellet volumes of STE buffer (10 mM Tris-HCl pH 8.0, 100 mM NaCl, 1 mM EDTA, protease inhibitors) containing lysozyme (0.1 mg/ml) were added to completely resuspend the pellets. After 15 min on ice, sarcosyl (N-lauryl sarcosine) was added to achieve a final concentration of 1.5% for protein denaturation. Sonication was performed using a tip probe sonicator (Branson Digital Sonifier 250) at 20% output or Bioruptor at max output, with 30s ON, 30s OFF pulses for 10 cycles, while keeping the sample on ice. Centrifugation was carried out at 10,000 xg, 4 °C for 10 min to remove insoluble material. The supernatant was saved, and Triton X-100 was added to achieve a final concentration of 5% for masking the sarcosyl and allowing protein refolding. This lysate was then added to glutathione (GSH) beaded agarose pre-equilibrated in STE buffer, and DTT was added to a final concentration of 2 mM. The mixture was rotated gently at 4° C for at least 4 h or overnight. Bead-binding reactions were spun at 500 xg for 5 min at 4° C. The supernatant was saved as the unbound fraction. MultiDSK agarose beads were washed twice with ice-cold STE containing 500 mM NaCl and 0.1% Triton X-100, followed by a wash in the same buffer with 50 mM NaCl. Two additional washes were performed with 1x PBS, and the beads were stored in 1x PBS with 0.02% sodium azide at 4 °C. MultiDSK agarose beads binding efficiency was assessed by boiling 25ul of beads in cracking buffer plus DTT and analyzing via SDS-PAGE. GST-MultiDSK protein runs at ∼60 kDa.

### MultiDSK pulldown

Packed MultiDSK agarose beads (25 μl) were employed to selectively deplete or enrich ubiquitylated proteins from a 1 mg total protein sample. Uniform volume adjustment was achieved by adding TENT buffer (50 mM Tris-HCl pH 7.4, 2 mM EDTA, 150 mM NaCl, 1% Triton X-100), which included protease inhibitors, phosphatase inhibitors, and freshly made 2 mM N-Ethylmaleimide. Final sample volumes ranged between 700 μl and 1 ml, with adjustments made for protein amounts as low as 500 μg, scaling both the volume of MultiDsk beads and the total reaction volume accordingly. Prewashing of beads was executed by spinning at 500 xg for 5 min at 4 °C. Following the removal of the supernatant, beads underwent a wash with TENT buffer containing protease inhibitors, phosphatase inhibitors, and 2 mM NEM. A well-resuspended MultiDsk bead slurry (typically 200 μl) was then aliquoted to each sample. The resulting mixture underwent rotation on a turning wheel/rotator (low to moderate speed) in the cold room for several hours to overnight. Samples were spun at 500 xg for 5 min at 4 °C, saving the supernatant as the “unbound” fraction. Beads were washed twice with 1 ml of TENT buffer containing protease inhibitors, phosphatase inhibitors, and 2 mM NEM, followed by a wash with 1x PBS containing the same inhibitors and NEM. The supernatant was removed, and to each bead sample, 40 μl of Laemmli buffer containing β-mercaptoethanol was added. The resulting supernatant, containing the enriched ubiquitylated proteins (referred to as the “bound” fraction), was prepared for WB analysis.

### ChIP-seq

Cells were cross-linked with ChIP Cross-link Gold (Diagenode, #C01019027) following manufacturer instructions. After incubation at room temperature for 30 min, the cells were fixed with a 1% formaldehyde solution (Thermo Fisher Scientific, #28908) for 10 min at room temperature. To stop the fixation, 125 mM glycine was added and incubated for 5 min. The cells were then washed twice with PBS and harvested on ice using cell scrapers. The harvested cells were resuspended in ChIP lysis buffer supplemented with protease inhibitors (20 mM Tris-HCl pH 8.0, 150 mM NaCl, 1 mM EDTA, 0.5 mM EGTA, 1% Triton X-100, 0.25% SDS) and sonicated on a Bioruptor Pico for 4 rounds of 10 cycles each at 4 °C with a short vortex between rounds to generate DNA fragments <500 bp. The sonicated cells were centrifuged, and the supernatants were transferred to new tubes and diluted with one volume of IP Dilution buffer (1% Triton X-100, 150 mM NaCl, 1 mM EDTA, 0.5 mM EGTA, 20 mM Tris-HCl pH 8.0, proteases and phosphatases inhibitors). For the immunoprecipitation step, 1 mg of pre-cleared protein extracts were incubated with 10 μg of antibodies, along with 1 μg of Spike-in antibody (Active Motif, #61686) and 50 ng of Spike-in chromatin from Drosophila (Active Motif, #53083) overnight at 4 °C with gentle rotation. For the negative control, Normal Rabbit IgG (Cell Signaling, #2729S) was employed. The next day, 50 μl of Protein A/G PLUS-Agarose beads (Santa Cruz, #sc-2003) were added to each IP reaction and incubated for 2 h at 4 °C with rotation. The beads were washed twice with IP Dilution buffer supplemented with 0.1% SDS (final), 3 times with ChIP wash buffer (IP Dilution buffer plus 0.5 M NaCl, 0.1% SDS, and 1% [w/v] Sodium Deoxycholate), and once with TE buffer (10mM Tris-HCl, 1mM EDTA) containing 50 mM NaCl. Elution of immunocomplexes from beads was performed with Elution Buffer (50 mM Tris-HCl, pH 8; 10 mM EDTA, 1% SDS) and incubation at 65 °C for 15 min. Then, the supernatants were reverse crosslinked at 65 °C for 3 h, treated sequentially with RNAse A (Sigma-Aldrich, #R5503) at 37 °C for 2 h, and then treated with Proteinase K (New England Biolabs, #P8107S) at 55 °C for 2 h. DNA purification was performed with the QIAquick PCR Purification Kit (Qiagen, #28106). Purified DNA was used to generate libraries using the NEBNext Ultra DNA Library Prep Kit for Illumina (New England Biolabs, #7370) following the manufacturer’s instructions. A Tapestation 2200 was used for fragment analysis using D1000 DNA ScreenTape (Agilent Technologies, #5067–5582). Libraries were quantified on a Qubit 4 fluorometer with Qubit double-stranded DNA high-sensitivity reagents (Thermo Fisher Scientific, #Q32851) following the manufacturer’s instructions, then pooled and sequenced (single or paired-end reads, 75 bp) on Illumina Novaseq 6000 (Oncogenomics Core Facility, SCCC, University of Miami).

For ChIP-seq experiments with DRB (5,6-Dichloro-1-β-d-ribofuranosylbenzimidazole), cells were treated with 100 μM DRB (Sigma-Aldrich, #D1916) 3 h before UV treatment (20 J/m^2^), and washed out 5 min after UV-irradiation. For ChIP-seq experiments where p300 or USP7 was inhibited, cells were treated with DMSO or A-485 (1 µM) 1 h before UV irradiation, and/or with FT671 (1 µM) 1 h after UV exposure (20 J/m²). For ChIP-seq experiments involving siRNA treatment, siRNA transfections were performed the day prior to the experiment, as previously described.

### ChIP-seq analyses

Single (or paired-end) read ChIP-seq fastq files were processed using the *ENCODE-DCC/chip-seq-pipeline2* pipeline (https://github.com/ENCODE-DCC/chip-seq-pipeline2) with default parameters and aligned to the hg19 genome. Peak calling was performed with *MACS2 v2.2.4*, and the peaks compared with each INPUT samples were used for subsequent analysis. *DeepTools v3.3.1* was used to generate the ChIP-seq peak profiles for each sample. Bigwig output files were visualized in the UCSC genome browser (https://genome.ucsc.edu). *HomerannotatePeaks v4.11* was used for peak annotation and gene assignment while *findMotifsGenome* was used for motif analysis. Pearson correlation between experimental replicates were calculated using *deepTools v3.3.1.* In the RPBII DRB-ChIP-seq analysis, BigWig files were filtered with IgG using *bigwigCompare* from *deepTools v3.5.3* with the following parameters: operation ratio, bin size (bs) 50, pseudocount 1, and skipNAs For strand-bias analysis, paired-end ChIPseq fastq files were aligned to the human genome hg19 using *BWA-MEM v 0.7.17-r1188*. The aligned files were converted to BAM files and sorted using *Samtools v1.9*. Duplicates reads were removed with *Picard v3.00*. Reads from each DNA strand (plus and minus) and split using *Samtools*. *bamCoverage v3.3.1* using RPKM option was used to generate normalized ChIP-seq peak profiles for each sample. The resulting BigWig output files were visualized in the UCSC Genome Browser (https://genome.ucsc.edu). The Strand Specificity Index (SSI) was calculated based on the numbers of forward and reverse mapped reads, using the formula presented in Figure 4F. Positive and negative SSI values indicate bidirectional gene orientation. Genes with fewer than 1 RPM were excluded from the analysis.

For RNAPII mapping to CPD dimers, the same files utilized in the strand bias analysis were normalized to RPCG (1x genome coverage) using *Deeptools v3.3.1*. Dimer sites and flanking regions, extending 400 bases from both ends, were extracted from genes longer than 20 kb on chromosome 1 of the hg19 genome sequence in the positive strand of DNA. Read coverage for each sample were calculated using computeMatrix from *Deeptools*. The results were visualized with plotHeatmap from *Deeptools*. The lowest value among the profiles for each sample was used for graph normalization.

### Chromatin-bound RNA (ChrRNA)-seq

Chromatin-bound RNA (ChrRNA)-seq method was conducted following the protocol described previously (Nojima et al., 2018). Briefly, 7x10^6^ cells for each condition were collected in ice-cold 1x PBS. Cells were pelleted at 400 xg for 5 min at 4 °C and then incubated in 4 mL of *Buffer 1* (10 mM Tris-HCl pH 8, 10 mM NaCl, 0.5% IGEPAL, and 2.5 mM MgCl_2_) for 5 min. Subsequently, 1 mL of ice-cold *Buffer 1* with the addition of 10% sucrose was under-layered, and nuclei were isolated by centrifugation for 5 min at 400 xg at 4 °C. The isolated nuclei were resuspended in 125 μl of *Buffer* 2 (20 mM Tris-HCl pH 8.0, 75 mM NaCl, 0.5 mM EDTA, 50% Glycerol) supplemented with 1x Complete EDTA-free protease inhibitors (Thermo Fisher Scientific, #A32955). The resuspended nuclei were then incubated on ice in 1.2 mL of *Buffer 3* (20 mM HEPES-KOH pH 7.6, 7.5 mM MgCl_2_, 0.2 mM EDTA, 300 mM NaCl, 1 M Urea, 1% IGEPAL) supplemented with 1x Complete EDTA-free protease inhibitors for 15 min. Chromatin-bound RNA was pelleted at 16,000 xg for 10 min at 4 °C, and the DNA was digested by incubating the chromatin pellet at 37 °C for 15 min shaking in 200 μ L *Buffer* 4 (10 mM Tris-HCl pH 8, 500 mM NaCl, and 10 mM MgCl_2_) with 0.25 U/μL TURBO DNase (Thermo Fisher Scientific, #AM2238). Proteins were digested by adding proteinase K solution (New England Biolabs, #P8107S) at 37 °C for 10 min. Chromatin-bound RNA was extracted with TRIzol Reagent (Thermo Fisher Scientific, #15596018) and isolated with Monarch Total RNA Miniprep kit (New England Biolabs, #T2010S) according to the manufacturer’s guidelines. The chromatin-bound RNA was dissolved in water, and RNA integrity was checked on the Agilent 4200 TapeStation system (Agilent Technologies). Chromatin-bound RNA (5 μg) was depleted of ribosomal RNA with RiboMinus Eukaryote System v2 (Thermo Fisher Scientific, #A15015) according to the manufacturer’s protocol. Libraries were prepared were prepared using NEBNext Ultra II RNA Library Prep Kit for Illumina (E7770) according to the manufacturer protocol. NEBNext Multiplex Oligos for Illumina (Unique Dual Index UMI Adaptors RNA Set 1, #E7416S/L) were used for the adapter ligation step. ERCC spike-in Mix 1 (Thermo Fisher Scientific, #4456740) was added to the samples for RNA-Seq normalization and sequencing was performed on a NovaSeq 6000 (Oncogenomics Core Facility, SCCC, University of Miami) with 150 bp paired-end reads.

### Polysome-enriched RNA-seq

For Polysome-enriched RNA purification, 15x10^6^ cells in two plates per condition and replicate were incubated with 1 mM 5-EU (MedChemExpress, HY-141140) for 20 min before the time-points 2 h and 3 h post-UV (20 J/m^2^) or non-irradiated (0 J/m^2^). Then, cells were incubated with 100 μg/ml cycloheximide (CHX) in growth media for 5 min at 37 °C and 5% CO_2_. Subsequently, cells were washed twice with 10 ml of ice-cold 1x PBS containing 100 μg/ml CHX. Cells were gently scraped, collected with 5 ml of 100 μg/ml PBS-CHX, and centrifuged at 300 xg for 5 min at 4 °C. Cells were resuspended in a *hypotonic buffer* (5 mM Tris-HCl pH 7.5, 2.5 mM MgCl_2_, 1.5 mM KCl and 1x EDTA-free protease inhibitor cocktail) with 100 μg/ml CHX, 1 μM DTT and 100 U of RNAse inhibitor (recombinant RNasin® Ribonuclease Inhibitor, Promega, #N2515). Sodium deoxycholate and Triton X-100 were added to the lysis reaction to a final concentration of 0.5%. After 10 minutes incubation at 4 °C, lysates were centrifuged at 16,000 xg for 7 min at 4 °C. The supernatant was transferred to a new pre-chilled 1.5 ml tube. Samples containing same OD_260 nm_ were added on top of a sucrose density gradient (from 50 to 5% sucrose, containing 20 mM HEPES (pH 7.6), 0.1 M KCl, 5 mM MgCl_2_, 10 µg/ml cycloheximide, 1x EDTA-free protease inhibitor and 10 units/ml RNase inhibitor) in Ultra-clear centrifuge tubes (Beckman Coulter, #344057). Samples were ultracentrifuged in a Sorvall WX 80+ (Thermo scientific, # 75000080) at 222,000 xg, for 2 h at 4 °C using an AH-650 rotor (Thermo-scientific, # 54294) without brakes. Fractions of 100 μl were collected and analyzed with a Nanodrop. Fractions corresponding to the polysomes were pooled and 1 μg of 5-EU labeled RNA of *Drosophila melanogaster* was added as a spike-in. Then, the samples were diluted in TRIzol LS Reagent (Thermo Fisher Scientific, #10296010). Purifications for WT or Δp300 polysomes were performed on different days.

RNA was purified according to the manufacturer protocol, using Monarch Total RNA purification columns (NEB, T2010S). The 5-EU RNA was purified using Click-iT™ Nascent RNA Capture Kit (ThermoFisher Scientific, C10365) according to manufacturer protocol. The RNA was depleted of ribosomal RNA with RiboMinus Eukaryote System v2 (Thermo Fisher Scientific, #A15015) according to the manufacturer’s protocol. RNA libraries were prepared using NEBNext Ultra II RNA Library Prep Kit for Illumina (NEB, E7770) according to the manufacturer protocol. Sequencing was performed on a NovaSeq X Plus (Oncogenomics Core Facility, SCCC, University of Miami) with 150 bp paired-end reads.

### TT-seq

Cells were incubated with 1 mM 517EU for 10 minutes and then collected using TRIzol LS Reagent (Thermo Fisher Scientific, #10296010). 517EU labeled RNA from *Drosophila melanogaster* was used as a spike17in. The RNA was purified using Monarch Total RNA Purification Columns (NEB, T2010S), and the 517EU labeled RNA was further purified with the Click17iT™ Nascent RNA Capture Kit (Thermo Fisher Scientific, C10365) according to the manufacturer’s protocol. Ribosomal RNA was depleted using the RiboMinus Eukaryote System v2 (Thermo Fisher Scientific, #A15015) following the manufacturer’s protocol. RNA libraries were prepared using the NEBNext Ultra II RNA Library Prep Kit for Illumina according to the manufacturer’s protocol. Sequencing was performed on a NovaSeq X Plus (Oncogenomics Core Facility, SCCC, University of Miami) with 150 bp paired17end reads.

### DRB and TT-seq

Samples were incubated with 100 μM DRB for 3 hours. Then, some cell groups were irradiated with 20 J/m², while others were left unirradiated (control groups). Immediately after UV irradiation, the cells were washed three times with warm PBS. Cells were incubated with DMEM 10% FBS and 2 mM 517EU was added 35 minutes after UV irradiation; 10 minutes later, the cells were harvested with TRIzol LS reagent (Invitrogen, 15596026). After purification, two protocols were followed: in one, the RNA was fragmented using the NEBNext Magnesium RNA Fragmentation Module (E610S), and in the other, the RNA was left intact. The 517EU RNA purification and library preparation were performed as described for the 5-EU RNAseq protocol.

### ChrRNA-seq analyses

RNA-seq FASTQ files were generated using *BCL converter v4.2.7* (Illumina) for index identification. Sample quality was analyzed with *FastQC v0.12.1*. Reads were trimmed using *Cutadapt v2.3* with the following parameters: -j 4 -m 18. Subsequently, reads were aligned to the hg19 genome using *STAR 2.7.10a* with the following parameters: chimSegmentMin 10, outSAMattributes All, outSAMstrandField intronMotif, alignIntronMax 1000000, alignEndsType EndToEnd, alignSJDBoverhangMin 6, quantMode TranscriptomeSAM, twopassMode basic. Duplicate reads were removed using *UMI-tools v1.1.4*. *FeatureCounts v2.0.3* was employed to calculate gene counts against either the human genome hg19 or the spike-in ERCC, using default parameters. Differential expression genes (DEGs) were identified using *DESeq2 v1.34.0* and R *v4.1.1*, setting a q-value threshold of less than 0.05 and log_2_FC of 1.5. The geometric mean of the ERCC spike-in was used for sample normalization. Volcano plots were generated using *ggplot2 v3.4.4.* GO enrichment analysis was performed using the WEB-based Gene Set Analysis Toolkit (WebGestalt) with the Molecular Function non-redundant database. GO terms with an FDR value of <0.05 were considered to be statistically significant and visualized using *ggplot2* R package.

For SNP and insertion-deletion mutation analysis, samples were aligned and analyzed using *Nextflow*. *RNAVar v1.0* (https://github.com/nf-core/rnavar) with default parameters, except for the STAR two-pass method and the feature_type gene for chromatin samples. Mutations identified in non-irradiated samples were used to clean the VCF files of UV-irradiated WT and Δp300 samples using *BCFtools v1.9*. The mutations were analyzed using *SnpEff v5.2* (Cingolani et al., 2012). Mutations intersecting with dimers in genes oriented in the positive direction on chromosome 1 were analyzed with SnpEff as previously indicated.

### ChrRNA-seq analyses

RNA-seq FASTQ files were generated using *BCL converter v4.2.7* (Illumina) for index identification. Sample quality was analyzed with *FastQC v0.12.1*. Reads were trimmed using *Cutadapt v2.3* with the following parameters: -j 4 -m 18. Subsequently, reads were aligned to the hg19 genome using *STAR 2.7.10a* with the following parameters: chimSegmentMin 10, outSAMattributes All, outSAMstrandField intronMotif, alignIntronMax 1000000, alignEndsType EndToEnd, alignSJDBoverhangMin 6, quantMode TranscriptomeSAM. Duplicate reads were removed using *UMI-tools v1.1.4*. *FeatureCounts v2.0.3* was employed to calculate gene counts against either the human genome hg19 or the spike-in ERCC, using default parameters. Differential expression genes (DEGs) were identified using *DESeq2 v1.34.0* and R *v4.1.1*, setting a q-value threshold of less than 0.05 and log_2_FC of 1.5. The geometric mean of the ERCC spike-in was used for sample normalization. Volcano plots were generated using *ggplot2 v3.4.4.* Similarities between experimental replicates were analyzed using *rlog* and *plotPCA* from the *DESeq2* package considering the top 10,000 genes. GO enrichment analysis was performed using the WEB-based Gene Set Analysis Toolkit (WebGestalt) with the Molecular Function non-redundant database. GO terms with an FDR value of <0.05 were considered to be statistically significant and visualized using *ggplot2* R package.

### Polysome and TT-seq analyses

FASTQ files were generated and processed as with ChrRNA-seq with a few changes. Briefly, reads were aligned to the human genome *hg19* and *D. melanogaster* genome using *STAR 2.7.10a*. Duplicate reads were removed using *Picard v3.00*. *FeatureCounts v2.0.3* was employed to calculate gene counts against either the human genome hg19 or the spike-in *D. melanogaster*, using default parameters. Differential expression genes (DEGs) were identified using *DESeq2 v1.34.0* and R *v4.1.1*, setting a q-value threshold of less than 0.05 and log_2_FC of 1.5. for Polysomes, and 1.0 for TT-seq. The geometric mean of the *D. melanogaster* spike-in was used for sample normalization. Volcano plots was performed as with ChrRNAseq. For splicing analysis, BAM files without duplicate reads were analyzed using *rMATS v4.2.0* with default options. Due to the random nature of UV damage, events with more than 4 reads in the inclusion counts were used for the analysis. Variants with more than 0.2 (variant gain) or less than -0.2 (variant loss) of inclusion level difference (ΔPSI), and an FDR of less than 0.05 were considered significant.

### DRB and TT-seq analyses

FASTQ files were processed and aligned as for Nascent RNA-seq. After the duplicated reads removal, samples were normalized according to their RPKM using *bamCoverage* from *deepTools v3.5.6.* A sample without 5-EU was used as negative control to filter the signal using *bigwigCompare* from *deepTools v3.5.6* with the following parameters: operation log2, bs 50, pseudocount 1 and skipNAs. Previous to generate the profile, samples were scaled to have the same background levels. The matrix was computed using genes with a size from 80 to 200 kbp from *hg19* genome, with a binsize of 1000 using *deepTools v3.5.6*. The profile was generated using *plotProfile* with the perGroup option.

### PRO-seq library preparation and data analysis

PRO-seq library preparation and data analysis followed established methodologies as outlined in previous studies(Beckedorff et al., 2020; Dasilva et al., 2021). Nuclear run-on assays were conducted using ten million nuclei, incubated for 3 minutes at 30 °C with 25 μM Biotin-11-ATP/UTP/CTP/GTP (PerkinElmer). Subsequently, total RNA was extracted and fragmented using 0.2 M NaOH for 10 minutes on ice, followed by purification of biotinylated nascent RNAs using streptavidin beads M-280 (Thermo Fisher Scientific, #11206D). Enzymatic steps involving RppH (NEB, #M0356S) and PNK (NEB, #M0201) were employed to remove the 5’ cap, repair the triphosphate, and repair the 5’ hydroxyl, respectively. Adapter ligation and reverse transcription using SuperScript III (Thermo Fisher Scientific, #56575) were performed, followed by PCR amplification. Size selection of libraries ranging from 140 to 350 bp was achieved using AMpure XP (Beckman Coulter, #A63882), and sequencing was conducted on a NovaSeq 6000 (Illumina) platform with single-read runs.

Raw fastq data underwent trimming using Cutadapt 1.14 and Trimmomatic v0.32 (Bolger et al., 2014; Martin, 2011) followed by alignment using bowtie 1.1.2 to either the hg19 or dm3 genome (Langmead et al., 2009). Strand-specific single nucleotide ends of aligned reads were constructed using BEDTools v2.28 with genomecov(Quinlan and Hall, 2010), and bedgraph data were normalized by the number of reads mapped to the spike-in dm3 genome before being transformed into bigwig data for further analysis. Read counts in the last 40% of the gene up to the transcription termination site (TTS) were calculated using FeatureCounts v2.0.3. Genes with differential polymerase activity were identified using *DESeq2 v1.34.0* and R *v4.1.1*, setting a q-value threshold of less than 0.05 and log2FC of 1.5. The geometric mean of the dm3 spike-in was used for sample normalization. Volcano plots were generated using *ggplot2 v3.4.*4. Similarities between experimental replicates were analyzed using *rlog* and *plotPCA* from the *DESeq2* package.

### Promoter Release Ratio analyses

Read counts in the promoter (-30 to +300 bp from TSS) or the gene body (first 20% of each gene from the promoter (+301 bp to 20%)) from PRO-seq data were obtained using *Bedtools v2.29.0*. Counts density were obtained by diving the read counts by region length. Promoter release ratio (PRR) for each gene was calculated as promoter read density divided by gene body read density. PRR values were ordered from minor to major values and the cumulative frequency was calculated for Δp300 and UV-irradiated Δp300.

### Cell cycle analysis

Cells (2x10^5^) were washed twice with cold 1x PBS and re-suspended in 1x PBS with 2 mM EDTA and fixed in 70 % ethanol overnight at 4 °C. Cells were recovered by centrifugation, washed with 1x PBS once and incubated at 37 °C for 30 min in staining buffer (2 mM EDTA in 1x PBS with 10 μg/ml RNase A, 5 μg/ml of propidium iodide (Sigma-Aldrich, # P4170). Cells were analyzed with CytoFLEX Flow Cytometer (Beckman Coulter) and FlowJo software (v10.7.1).

### Apoptosis assay

For the staining process, cells were initially washed twice with cold PBS-BSA 0.5%, followed by resuspension in 1x Binding Buffer (10 mM HEPES pH 7.4, 140 mM NaCl, 2,5 mM CaCl_2_) at a concentration of 1x10^6^ cells/ml. Subsequently, 100 μl of this solution was transferred to a 5 ml culture tube. APC Annexin V (5 μl, BioLegend, #640941) and propidium iodide (10 μl of a 50 μg/ml solution) were added, and the cells were gently vortexed before incubating for 15 min at RT in darkness. Finally, 400 μl of 1x Binding Buffer was added to each tube, and the samples were analyzed by flow cytometry (CytoFLEX, Beckman Coulter) within 30 min. Cells staining positive for APC Annexin V and negative for PI indicate cells in the early stages of apoptosis. Cells positive for both APC Annexin V and PI may be in the end stage of apoptosis, undergoing necrosis, or already dead. Negative staining for both APC Annexin V and PI suggests live cells not undergoing measurable apoptosis.

### Global Nascent 5-EU RNA labeling

HeLa, U2OS, or IMR90 cells were grown to 50% confluence on glass coverslips and were either exposed to 5 J/m^2^ UV or not exposed. DMSO or A-485 (1 μM) were added 1 h before UV exposure. In all cases, 1 mM 5-ethynyluridine (5’-EU) was added 20 min before collection at which point cells were fixated with 3.7% paraformaldehyde for 15 min at RT. After two washes with 1x PBS and cell permeabilization with 0.5% Triton X-100 in 1x PBS for 15 min, 5’-EU labelling was carried using a Click-iT RNA Alexa Fluor 488 Imaging kit (Thermo Fisher Scientific, #C10329) following manufacturer instructions. After Click-iT reaction, cells were incubated with 0.5 μg/ml, 6-diamino-2-phenylindole (DAPI) in 1x PBS for 5 min, washed with 1x PBS for 2 min, air dried, and mounted in SlowFade Diamond antifade mounting reagent (Thermo Fisher Scientific, #S36963). Samples were analyzed using a Leica DMI6000B microscope with LASX software (Leica).

### Unscheduled DNA synthesis (UDS)

UDS was visualized and quantified as previously described (van der Meer et al., 2023). Briefly, HeLa cells were transfected with siRNA targeting XPC to inhibit global genome nucleotide excision repair (GG-NER). Following transfection, cells were subjected to serum starvation in DMEM without fetal bovine serum for 24 hours. To induce DNA damage, cells were locally UV-irradiated through 5 mm pore filters (Millipore, TMTP04700) at a dose of 100 J/m² for TC-NER UDS. Immediately after irradiation, cells were pulse-labeled with 50 μM 5-ethynyl-2’-deoxyuridine (EdU; MCE, #HY-118411) and 0.5 μM 5-fluoro-2-deoxyuridine (Santa Cruz, # sc-202425).

After labeling, cells were fixed in 2% paraformaldehyde in 1x PBS for 10 minutes at room temperature and stored in 1x PBS. Permeabilization was performed by incubating cells for 10 minutes in Blocking Solution (1% BSA, 3% horse serum, 0.2 % Triton X-100 in 1x PBS) at room temperature. EdU incorporation was then detected via click chemistry by incubating cells for 1 hour in a reaction mixture containing 6 μM AZDYE 488 azide (Vector Labs, #CCT-1275-5), 10 mM ascorbic acid, 4 mM copper sulfate, and 50 mM Tris-HCl buffer pH 8.0. Following EdU detection, cells were fixed with 2% paraformaldehyde in 1x PBS for 10 minutes, and then wash three times with 1x PBS.

To visualize UV-induced DNA damage, cells were extensively washed with PBS, and DNA was denatured using 0.5 N NaOH for 10 minutes. Blocking was performed using blocking buffer supplemented with 2% horse serum for 30 minutes. Cells were then incubated for 2 hours with mouse anti-CPD antibody (Cosmo Bio, # NMDND001) in blocking buffer, followed by five washes with 1x PBS (5 minutes each). Cells were then stained with goat anti-mouse Alexa 594 in permeabilization buffer for 30 minutes, washed three times with 1x PBS, counterstained with 0.2 mg/ml DAPI, washed again with 1x PBS, rinsed with Milli-Q water, and air-dried. Finally, samples were mounted in SlowFade Diamond antifade mounting reagent (ThermoFisher Sc.) and analyzed using a Leica DMI6000B microscope with LASX software (Leica).

### Immunofluorescence assays

Cells were fixed with 2% paraformaldehyde in 1x PBS for 10 min at room temperature (RT), washed with 1x PBS and bound to poly-L-lysine coated slides. After wash with 1x PBS for 5 min, cells were incubated with Blocking Solution (1 mg/ml BSA, 3% goat serum, 0.2 % Triton X-100 in 1x PBS) for 30 min at RT. Next, the cells were incubated with the primary antibodies in Blocking Solution for 1 h at RT, washed 3 times with 1x PBS, and incubated with Alexa 488 or 594 coupled secondary antibodies (Thermo Fisher Scientific) for another 30 min. Finally, cells were incubated with 0.5 μg/ml, 6-diamino-2-phenylindole (DAPI) in 1x PBS for 5 min, washed with 1x PBS for 2 min, air dried, and mounted in SlowFade Diamond antifade mounting reagent (Thermo Fisher Scientific, #S36963). Samples were analyzed using a Leica DMI6000B microscope with LASX software (Leica).

## References

Adam, S., Polo, S.E., and Almouzni, G. (2013). Transcription recovery after DNA damage requires chromatin priming by the H3.3 histone chaperone HIRA. Cell 155, 94–106.

Agapov, A., Olina, A., and Kulbachinskiy, A. (2022). RNA polymerase pausing, stalling and bypass during transcription of damaged DNA: from molecular basis to functional consequences. Nucleic Acids Res 50, 3018–3041.

Anindya, R., Aygun, O., and Svejstrup, J.Q. (2007). Damage-induced ubiquitylation of human RNA polymerase II by the ubiquitin ligase Nedd4, but not Cockayne syndrome proteins or BRCA1. Mol Cell 28, 386–397.

Bannister, A.J., and Kouzarides, T. (1996). The CBP co-activator is a histone acetyltransferase. Nature 384, 641–643.

Beckedorff, F., Blumenthal, E., daSilva, L.F., Aoi, Y., Cingaram, P.R., Yue, J., Zhang, A., Dokaneheifard, S., Valencia, M.G., Gaidosh, G., et al. (2020). The Human Integrator Complex Facilitates Transcriptional Elongation by Endonucleolytic Cleavage of Nascent Transcripts. Cell reports 32, 107917.

Bolger, A.M., Lohse, M., and Usadel, B. (2014). Trimmomatic: a flexible trimmer for Illumina sequence data. Bioinformatics 30, 2114–2120.

Bregman, D.B., Halaban, R., van Gool, A.J., Henning, K.A., Friedberg, E.C., and Warren, S.L. (1996). UV-induced ubiquitination of RNA polymerase II: a novel modification deficient in Cockayne syndrome cells. Proc Natl Acad Sci U S A 93, 11586–11590.

Brueckner, F., Hennecke, U., Carell, T., and Cramer, P. (2007). CPD damage recognition by transcribing RNA polymerase II. Science 315, 859–862.

Cingolani, P., Platts, A., Wang le, L., Coon, M., Nguyen, T., Wang, L., Land, S.J., Lu, X., and Ruden, D.M. (2012). A program for annotating and predicting the effects of single nucleotide polymorphisms, SnpEff: SNPs in the genome of Drosophila melanogaster strain w1118; iso-2; iso-3. Fly (Austin) 6, 80–92.

Dancy, B.M., and Cole, P.A. (2015). Protein lysine acetylation by p300/CBP. Chem Rev 115, 2419–2452.

Dasilva, L.F., Blumenthal, E., Beckedorff, F., Cingaram, P.R., Gomes Dos Santos, H., Edupuganti, R.R., Zhang, A., Dokaneheifard, S., Aoi, Y., Yue, J., et al. (2021). Integrator enforces the fidelity of transcriptional termination at protein-coding genes. Sci Adv 7, eabe3393.

Doetsch, P.W. (2002). Translesion synthesis by RNA polymerases: occurrence and biological implications for transcriptional mutagenesis. Mutat Res 510, 131–140.

Epanchintsev, A., Costanzo, F., Rauschendorf, M.A., Caputo, M., Ye, T., Donnio, L.M., Proietti-de-Santis, L., Coin, F., Laugel, V., and Egly, J.M. (2017). Cockayne’s Syndrome A and B Proteins Regulate Transcription Arrest after Genotoxic Stress by Promoting ATF3 Degradation. Mol Cell 68, 1054–1066 e1056.

Fei, J., and Chen, J. (2012). KIAA1530 protein is recruited by Cockayne syndrome complementation group protein A (CSA) to participate in transcription-coupled repair (TCR). J Biol Chem 287, 35118–35126.

Gregersen, L.H., and Svejstrup, J.Q. (2018). The Cellular Response to Transcription-Blocking DNA Damage. Trends Biochem Sci 43, 327–341.

Groisman, R., Kuraoka, I., Chevallier, O., Gaye, N., Magnaldo, T., Tanaka, K., Kisselev, A.F., Harel-Bellan, A., and Nakatani, Y. (2006). CSA-dependent degradation of CSB by the ubiquitin-proteasome pathway establishes a link between complementation factors of the Cockayne syndrome. Genes Dev 20, 1429–1434.

Gyenis, A., Umlauf, D., Ujfaludi, Z., Boros, I., Ye, T., and Tora, L. (2014). UVB induces a genome-wide acting negative regulatory mechanism that operates at the level of transcription initiation in human cells. PLoS Genet 10, e1004483.

Henning, K.A., Li, L., Iyer, N., McDaniel, L.D., Reagan, M.S., Legerski, R., Schultz, R.A., Stefanini, M., Lehmann, A.R., Mayne, L.V., et al. (1995). The Cockayne syndrome group A gene encodes a WD repeat protein that interacts with CSB protein and a subunit of RNA polymerase II TFIIH. Cell 82, 555–564.

Higa, M., Tanaka, K., and Saijo, M. (2018). Inhibition of UVSSA ubiquitination suppresses transcription-coupled nucleotide excision repair deficiency caused by dissociation from USP7. FEBS J 285, 965–976.

Kelland, L. (2007). The resurgence of platinum-based cancer chemotherapy. Nat Rev Cancer 7, 573–584.

Kleiman, F.E., Wu-Baer, F., Fonseca, D., Kaneko, S., Baer, R., and Manley, J.L. (2005). BRCA1/BARD1 inhibition of mRNA 3’ processing involves targeted degradation of RNA polymerase II. Genes Dev 19, 1227–1237.

Kokic, G., Wagner, F.R., Chernev, A., Urlaub, H., and Cramer, P. (2021). Structural basis of human transcription-DNA repair coupling. Nature 598, 368–372.

Kokic, G., Yakoub, G., van den Heuvel, D., Wondergem, A.P., van der Meer, P.J., van der Weegen, Y., Chernev, A., Fianu, I., Fokkens, T.J., Lorenz, S., et al. (2024). Structural basis for RNA polymerase II ubiquitylation and inactivation in transcription-coupled repair. Nat Struct Mol Biol.

Kwak, H., Fuda, N.J., Core, L.J., and Lis, J.T. (2013). Precise maps of RNA polymerase reveal how promoters direct initiation and pausing. Science 339, 950–953.

Langmead, B., Trapnell, C., Pop, M., and Salzberg, S.L. (2009). Ultrafast and memory-efficient alignment of short DNA sequences to the human genome. Genome Biol 10, R25.

Lans, H., Hoeijmakers, J.H.J., Vermeulen, W., and Marteijn, J.A. (2019). The DNA damage response to transcription stress. Nat Rev Mol Cell Biol 20, 766–784.

Lasko, L.M., Jakob, C.G., Edalji, R.P., Qiu, W., Montgomery, D., Digiammarino, E.L., Hansen, T.M., Risi, R.M., Frey, R., Manaves, V., et al. (2017). Discovery of a selective catalytic p300/CBP inhibitor that targets lineage-specific tumours. Nature 550, 128–132.

Lavigne, M.D., Konstantopoulos, D., Ntakou-Zamplara, K.Z., Liakos, A., and Fousteri, M. (2017). Global unleashing of transcription elongation waves in response to genotoxic stress restricts somatic mutation rate. Nature communications 8, 2076.

Li, W., Selvam, K., Ko, T., and Li, S. (2014). Transcription bypass of DNA lesions enhances cell survival but attenuates transcription coupled DNA repair. Nucleic Acids Res 42, 13242–13253.

Liebelt, F., Schimmel, J., Verlaan-de Vries, M., Klemann, E., van Royen, M.E., van der Weegen, Y., Luijsterburg, M.S., Mullenders, L.H., Pines, A., Vermeulen, W., et al. (2020). Transcription-coupled nucleotide excision repair is coordinated by ubiquitin and SUMO in response to ultraviolet irradiation. Nucleic Acids Res 48, 231–248.

Martin, M. (2011). Cutadapt Removes Adapter Sequences From High-Throughput Sequencing Reads. EMBnet Journal 17, 10–12.

Mayne, L.V., and Lehmann, A.R. (1982). Failure of RNA synthesis to recover after UV irradiation: an early defect in cells from individuals with Cockayne’s syndrome and xeroderma pigmentosum. Cancer Res 42, 1473–1478.

Nakazawa, Y., Hara, Y., Oka, Y., Komine, O., van den Heuvel, D., Guo, C., Daigaku, Y., Isono, M., He, Y., Shimada, M., et al. (2020). Ubiquitination of DNA Damage-Stalled RNAPII Promotes Transcription-Coupled Repair. Cell 180, 1228–1244 e1224.

Nakazawa, Y., Sasaki, K., Mitsutake, N., Matsuse, M., Shimada, M., Nardo, T., Takahashi, Y., Ohyama, K., Ito, K., Mishima, H., et al. (2012). Mutations in UVSSA cause UV-sensitive syndrome and impair RNA polymerase IIo processing in transcription-coupled nucleotide-excision repair. Nat Genet 44, 586–592.

Nieto Moreno, N., Olthof, A.M., and Svejstrup, J.Q. (2023). Transcription-Coupled Nucleotide Excision Repair and the Transcriptional Response to UV-Induced DNA Damage. Annu Rev Biochem 92, 81–113.

Noe Gonzalez, M., Blears, D., and Svejstrup, J.Q. (2021). Causes and consequences of RNA polymerase II stalling during transcript elongation. Nat Rev Mol Cell Biol 22, 3–21.

Nojima, T., Tellier, M., Foxwell, J., Ribeiro de Almeida, C., Tan-Wong, S.M., Dhir, S., Dujardin, G., Dhir, A., Murphy, S., and Proudfoot, N.J. (2018). Deregulated Expression of Mammalian lncRNA through Loss of SPT6 Induces R-Loop Formation, Replication Stress, and Cellular Senescence. Mol Cell 72, 970–984 e977.

Ogryzko, V.V., Schiltz, R.L., Russanova, V., Howard, B.H., and Nakatani, Y. (1996). The transcriptional coactivators p300 and CBP are histone acetyltransferases. Cell 87, 953–959.

Okuda, M., Nakazawa, Y., Guo, C., Ogi, T., and Nishimura, Y. (2017). Common TFIIH recruitment mechanism in global genome and transcription-coupled repair subpathways. Nucleic Acids Res 45, 13043–13055.

Proietti-De-Santis, L., Drane, P., and Egly, J.M. (2006). Cockayne syndrome B protein regulates the transcriptional program after UV irradiation. EMBO J 25, 1915–1923.

Quinlan, A.R., and Hall, I.M. (2010). BEDTools: a flexible suite of utilities for comparing genomic features. Bioinformatics 26, 841–842.

Ratner, J.N., Balasubramanian, B., Corden, J., Warren, S.L., and Bregman, D.B. (1998). Ultraviolet radiation-induced ubiquitination and proteasomal degradation of the large subunit of RNA polymerase II. Implications for transcription-coupled DNA repair. J Biol Chem 273, 5184–5189.

Rockx, D.A., Mason, R., van Hoffen, A., Barton, M.C., Citterio, E., Bregman, D.B., van Zeeland, A.A., Vrieling, H., and Mullenders, L.H. (2000). UV-induced inhibition of transcription involves repression of transcription initiation and phosphorylation of RNA polymerase II. Proc Natl Acad Sci U S A 97, 10503–10508.

Schwertman, P., Lagarou, A., Dekkers, D.H., Raams, A., van der Hoek, A.C., Laffeber, C., Hoeijmakers, J.H., Demmers, J.A., Fousteri, M., Vermeulen, W., et al. (2012). UV-sensitive syndrome protein UVSSA recruits USP7 to regulate transcription-coupled repair. Nat Genet 44, 598–602.

Starita, L.M., Horwitz, A.A., Keogh, M.C., Ishioka, C., Parvin, J.D., and Chiba, N. (2005). BRCA1/BARD1 ubiquitinate phosphorylated RNA polymerase II. J Biol Chem 280, 24498–24505.

Troelstra, C., van Gool, A., de Wit, J., Vermeulen, W., Bootsma, D., and Hoeijmakers, J.H. (1992). ERCC6, a member of a subfamily of putative helicases, is involved in Cockayne’s syndrome and preferential repair of active genes. Cell 71, 939–953.

Tufegdzic Vidakovic, A., Mitter, R., Kelly, G.P., Neumann, M., Harreman, M., Rodriguez-Martinez, M., Herlihy, A., Weems, J.C., Boeing, S., Encheva, V., et al. (2020). Regulation of the RNAPII Pool Is Integral to the DNA Damage Response. Cell 180, 1245–1261 e1221.

Turnbull, A.P., Ioannidis, S., Krajewski, W.W., Pinto-Fernandez, A., Heride, C., Martin, A.C.L., Tonkin, L.M., Townsend, E.C., Buker, S.M., Lancia, D.R., et al. (2017). Molecular basis of USP7 inhibition by selective small-molecule inhibitors. Nature 550, 481–486.

van den Boom, V., Citterio, E., Hoogstraten, D., Zotter, A., Egly, J.M., van Cappellen, W.A., Hoeijmakers, J.H., Houtsmuller, A.B., and Vermeulen, W. (2004). DNA damage stabilizes interaction of CSB with the transcription elongation machinery. J Cell Biol 166, 27–36.

van der Meer, P.J., Van Den Heuvel, D., and Luijsterburg, M.S. (2023). Unscheduled DNA Synthesis at Sites of Local UV-induced DNA Damage to Quantify Global Genome Nucleotide Excision Repair Activity in Human Cells. Bio Protoc 13.

van der Weegen, Y., de Lint, K., van den Heuvel, D., Nakazawa, Y., Mevissen, T.E.T., van Schie, J.J.M., San Martin Alonso, M., Boer, D.E.C., Gonzalez-Prieto, R., Narayanan, I.V., et al. (2021). ELOF1 is a transcription-coupled DNA repair factor that directs RNA polymerase II ubiquitylation. Nat Cell Biol 23, 595–607.

van der Weegen, Y., Golan-Berman, H., Mevissen, T.E.T., Apelt, K., Gonzalez-Prieto, R., Goedhart, J., Heilbrun, E.E., Vertegaal, A.C.O., van den Heuvel, D., Walter, J.C., et al. (2020). The cooperative action of CSB, CSA, and UVSSA target TFIIH to DNA damage-stalled RNA polymerase II. Nature communications 11, 2104.

van Gool, A.J., Citterio, E., Rademakers, S., van Os, R., Vermeulen, W., Constantinou, A., Egly, J.M., Bootsma, D., and Hoeijmakers, J.H. (1997). The Cockayne syndrome B protein, involved in transcription-coupled DNA repair, resides in an RNA polymerase II-containing complex. EMBO J 16, 5955–5965.

Vermeulen, W., and Fousteri, M. (2013). Mammalian transcription-coupled excision repair. Cold Spring Harbor perspectives in biology 5, a012625.

Walmacq, C., Cheung, A.C., Kireeva, M.L., Lubkowska, L., Ye, C., Gotte, D., Strathern, J.N., Carell, T., Cramer, P., and Kashlev, M. (2012). Mechanism of translesion transcription by RNA polymerase II and its role in cellular resistance to DNA damage. Mol Cell 46, 18–29.

Wang, W., Walmacq, C., Chong, J., Kashlev, M., and Wang, D. (2018). Structural basis of transcriptional stalling and bypass of abasic DNA lesion by RNA polymerase II. Proc Natl Acad Sci U S A 115, E2538–E2545.

Williamson, L., Saponaro, M., Boeing, S., East, P., Mitter, R., Kantidakis, T., Kelly, G.P., Lobley, A., Walker, J., Spencer-Dene, B., et al. (2017). UV Irradiation Induces a Non-coding RNA that Functionally Opposes the Protein Encoded by the Same Gene. Cell 168, 843–855 e813.

Wilson, M.D., Saponaro, M., Leidl, M.A., and Svejstrup, J.Q. (2012). MultiDsk: a ubiquitin-specific affinity resin. PloS one 7, e46398.

Xu, J., Lahiri, I., Wang, W., Wier, A., Cianfrocco, M.A., Chong, J., Hare, A.A., Dervan, P.B., DiMaio, F., Leschziner, A.E., et al. (2017). Structural basis for the initiation of eukaryotic transcription-coupled DNA repair. Nature 551, 653–657.

Yasukawa, T., Kamura, T., Kitajima, S., Conaway, R.C., Conaway, J.W., and Aso, T. (2008). Mammalian Elongin A complex mediates DNA-damage-induced ubiquitylation and degradation of Rpb1. EMBO J 27, 3256–3266.

Zhang, C., Xu, C., Gao, X., and Yao, Q. (2022). Platinum-based drugs for cancer therapy and anti-tumor strategies. Theranostics 12, 2115–2132.

Zhang, X., Horibata, K., Saijo, M., Ishigami, C., Ukai, A., Kanno, S., Tahara, H., Neilan, E.G., Honma, M., Nohmi, T., et al. (2012). Mutations in UVSSA cause UV-sensitive syndrome and destabilize ERCC6 in transcription-coupled DNA repair. Nat Genet 44, 593–597.

