## Supplementary material for "Bypass of Blocking Lesions by RNAPII Impairs the Transcriptional DNA Damage Response": S1

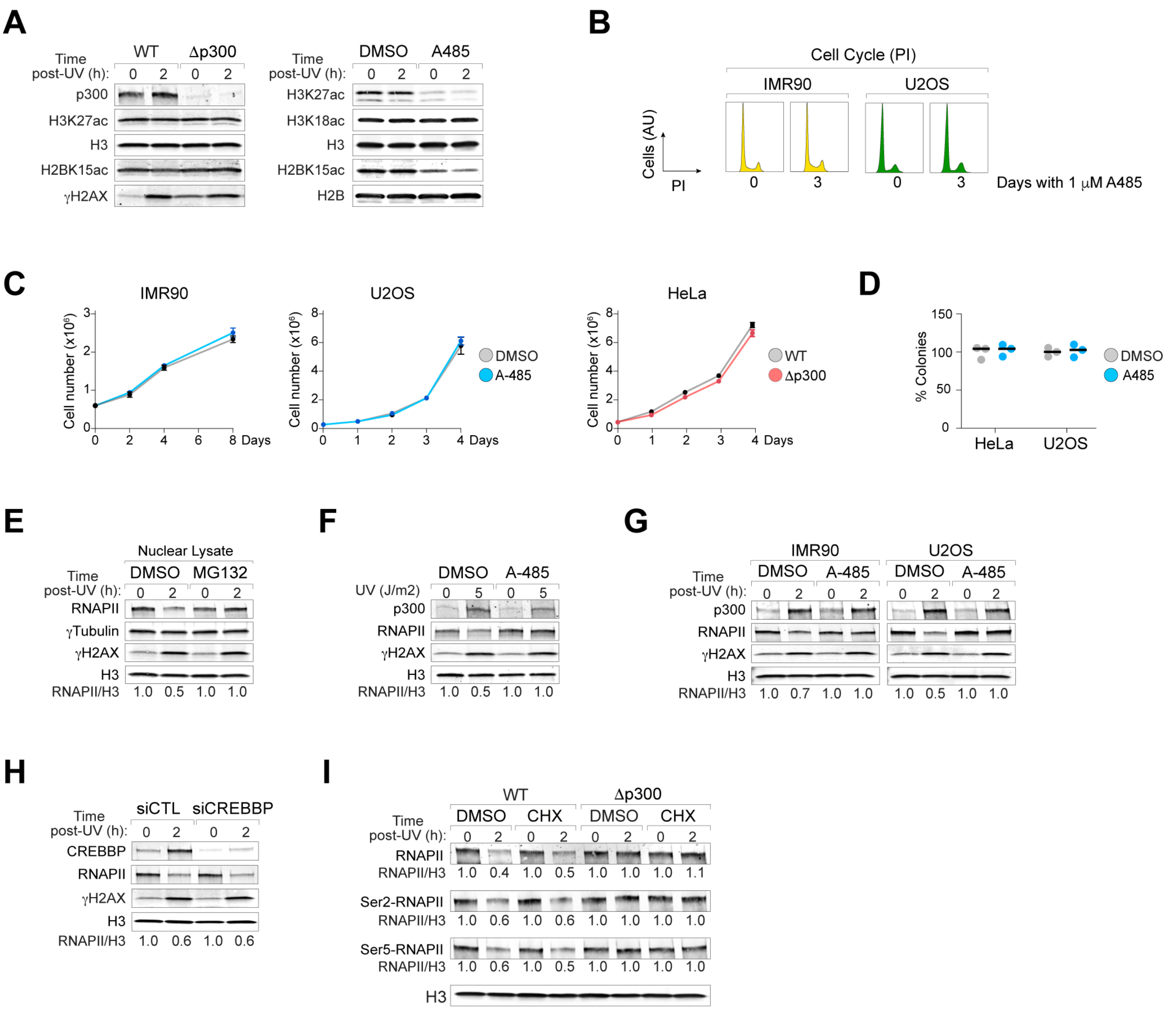


**Figure S1. Related to Figure 1. (A)** WB analysis of histone extracts from WT or Δp300 HeLa cells (left) and in the presence of DMSO or 1 μM A-485 one hour prior to UV exposure (20 J/m$^2$). **(B)** Propidium iodide (PI) staining of IMR90 and U2OS cells exposed to DMSO or 1 μM A-485 for 3 days. **(C)** *Left*, growth curves for IMR90 and U2OS cells in presence of DMSO or 1 μM A-485 during 8 or 4 days respectively. *Right,* growth curves for HeLa WT or Δp300 cells for 4 days. **(D)** Percentage of colonies formed by HeLa and U2OS cells exposed to either DMSO or 1 μM A-485 over 10 days. **(E)** WB analysis of nuclear extracts from HeLa cells treated with 10 μM MG132 (proteasome inhibitor), collected at 0 and 2 h after UV exposure (20 J/m$^2$). The ratios of RNAPII and H3 signals relative to samples before UV damage are shown. **(F)** WB analysis of chromatin-enriched extracts from HeLa cells collected 2 h post-UV (5 J/m$^2$). The ratios of RNAPII and H3 signals relative to samples before UV damage are shown. **(G)** WB analysis of chromatin-enriched extracts from IMR90 and U2OS cells collected at 0 or 2 h post-UV (20 J/m$^2$). The ratios of RNAPII and H3 signals relative to samples before UV damage are shown. **(H)** WB analysis of the indicated proteins in chromatin-enriched fractions from HeLa cells transfected with siCTL or siCREBBP. Cells were collected 2 h post-UV exposure (20 J/$m^2$). The ratios of RNAPII and H3 signals relative to samples before UV damage are shown **(I)** WB analysis of chromatin-enriched fractions from HeLa WT and ∆p300 incubated with DMSO or CHX and collected 2 h post-UV exposure (20 J/$m^2$). The ratios of RNAPII (total, Ser2, Ser5) and H3 signals relative to samples before UV damage are shown. All experiments were performed in at least three independent experimental replicates
