## Supplementary material for "Bypass of Blocking Lesions by RNAPII Impairs the Transcriptional DNA Damage Response": S2

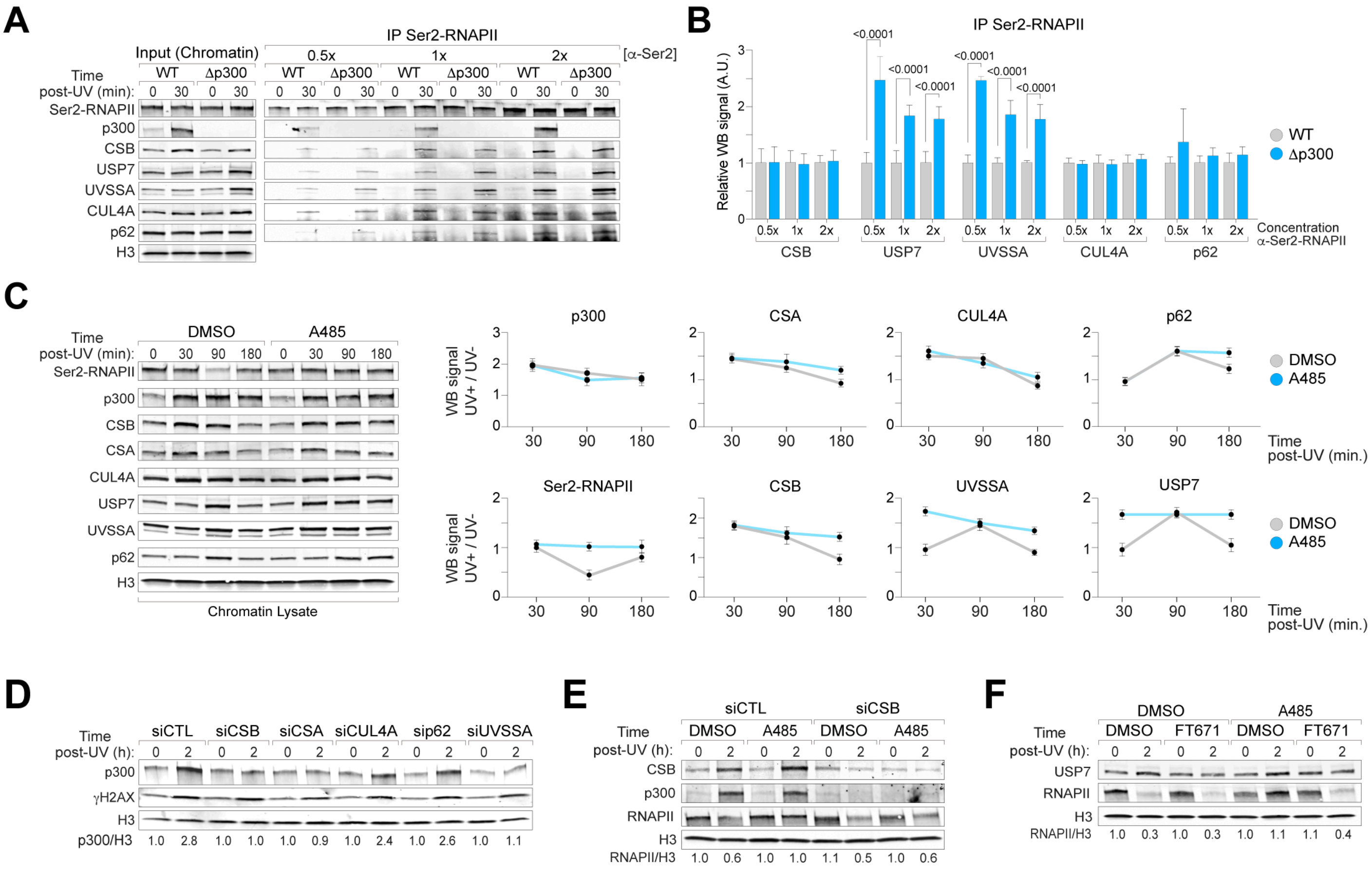


**Figure S2. Related to Figure 2. (A)** WB analyses of the indicated proteins in IP assays using UVSSA or USP7 antibodies on samples collected 30 min post-UV irradiation (20 J/m$^2$) from chromatin-enriched extracts of HeLa cells. DMSO or 1 μM A-485 was added 1 h before UV. The input represents 3% of the chromatin fraction. **(B)** WB analyses for the indicated proteins of IP assays using p300 or UVSSA antibodies on samples collected at 0 and 30 min post-UV exposure (20 J/m$^2$) from chromatin-enriched extracts of HeLa cells. DMSO or 1 μM A-485 was added 1 h before UV-irradiation. The input represents 3% of the chromatin fraction. **(C)** WB analysis showing the efficiency of gene knockdown by different siRNAs before and after UV exposure (20 J/m$^2$). **(D)** WB analyses from WT or ΔUSP7 HeLa cells at 0 and 2 h post-UV (20 J/m$^2$). DMSO or 1 μM A-485 was added 1 h prior to UV exposure. The ratios of RNAPII and H3 signals relative to samples before UV damage are shown. All experiments were performed in at least three independent experimental replicates.
