## Supplementary material for "Bypass of Blocking Lesions by RNAPII Impairs the Transcriptional DNA Damage Response": S3

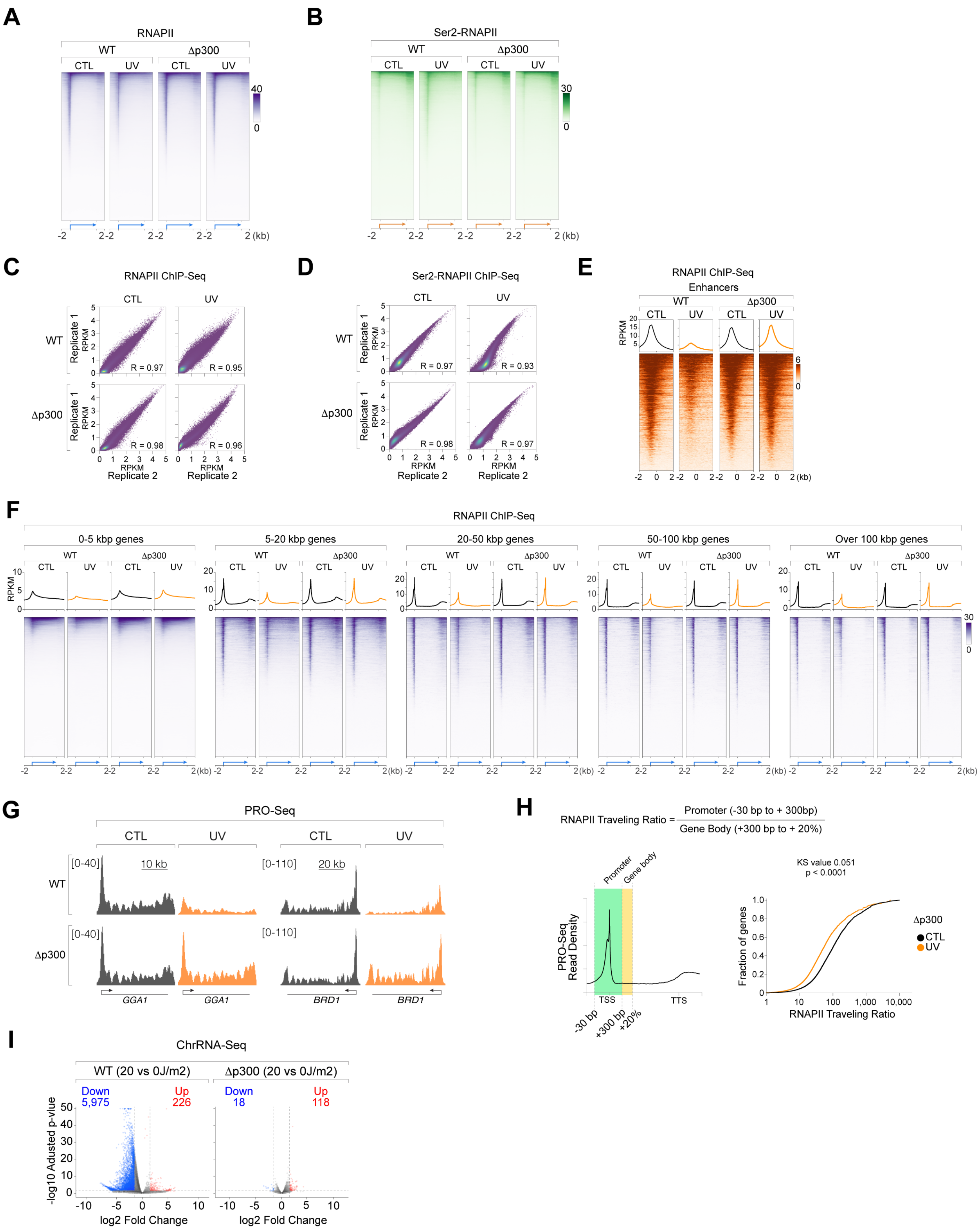


**Figure S3. Related to Figure 3. (A)** ChIP-seq heatmaps of total RNAPII from *Fig. 3A*. Rows are sorted in decreasing order by RNAPII occupancy in the indicated region. Color-scaled intensities are in units of rpm. **(B)** As in *A* but for Ser2-RNAPII from *Fig. 3C*. **(C)** Pearson correlation between normalized mapped reads from experimental replicates of RNAPII ChIP-seq. **(D)** As in *C* but for Ser2-RNAPII ChIP-seq. **(E)** Distribution of chromatin-bound RNAPII within gene bodies and flanking regions before and 2 h after UV irradiation (20 $J/m^2$) in the enhancer regions of both WT and Δp300 HeLa cells (RPKM, Reads Per Kilobase of transcript per Million mapped reads). **(F)** ChIP-seq heatmap of total RNAPII for different gene size clusters (0-5; 5-20; 20-50; 50-100; over-100 kilobase pair). **(G)** Track examples of the PRO-seq from *Fig 3F*. **(H)** *Left*, schematic representation of the modified traveling ratio (promoter/gene body density ratio), used to calculate the active RNAPII release ratio (PRR). The promoter is defined as the region covering 30 bp upstream to 300 bp downstream of the TSS; the gene body is defined as the region from 300 bp to 20% of the gene body downstream of the TSS. PRO-seq read counts were used for the analysis. *Right*, cumulative distribution plot of the PRR distribution in Δp300 HeLa cells before and after UV-irradiation. Genes with a promoter density less than 0.1 were excluded. A Kolmogorov-Smirnov test was used to determine the significance of the differences between the two curves. **(I)** Volcano plots of differential expressed genes ($log_2FC \geq 1.5$, q-value < 0.05) derived from chromatin-associated RNA-seq analyses comparing 20 $J/m^2$ vs 0 $J/m^2$ in HeLa WT or Δp300 cells. Samples were normalized according to their spike-in geometric mean. The number of downregulated genes is shown in blue, and the number of upregulated genes is shown in red. All experiments were performed in at least two independent experimental replicates.
