## Supplementary material for "Bypass of Blocking Lesions by RNAPII Impairs the Transcriptional DNA Damage Response": S4

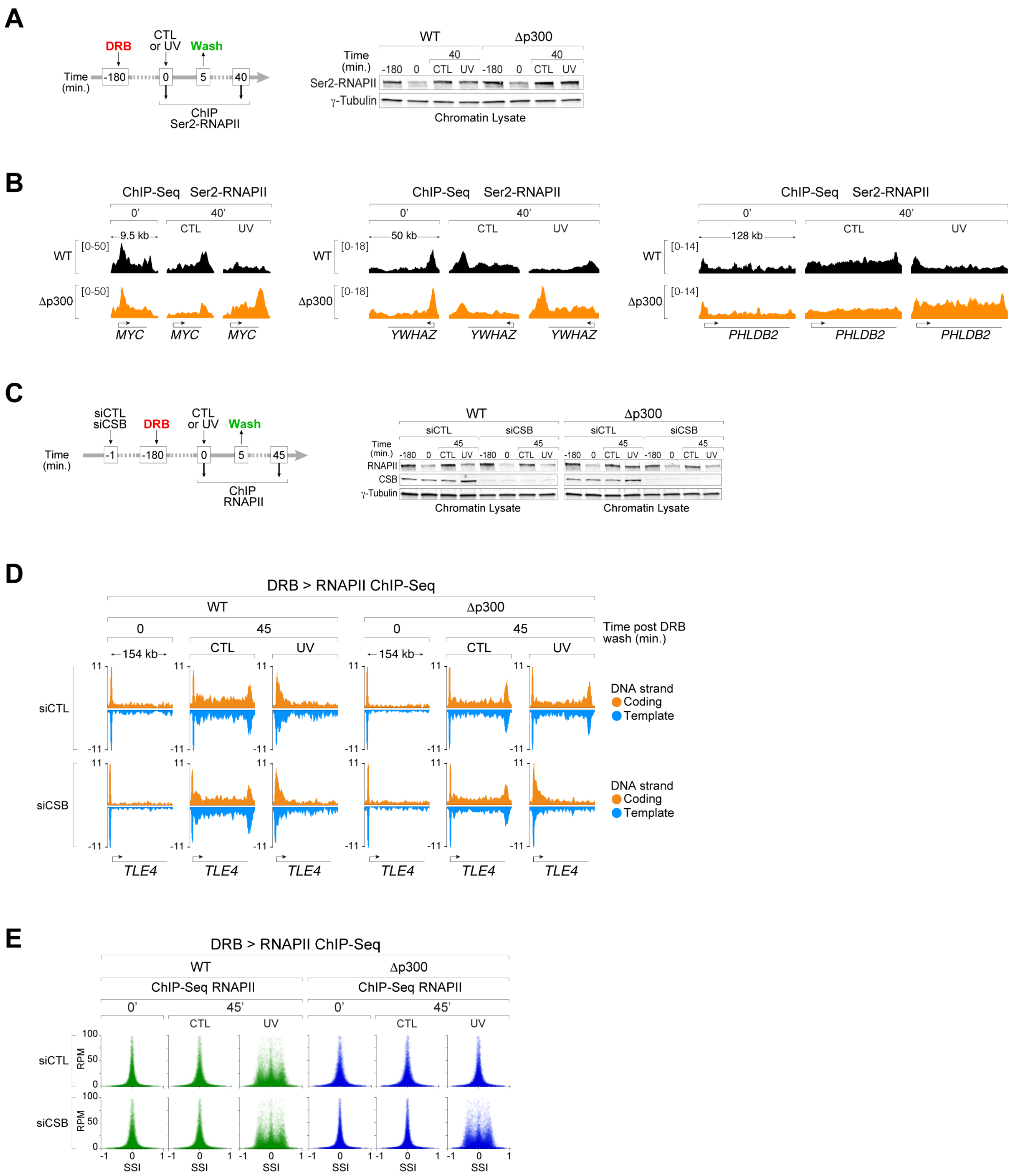


**Figure S4. Related to Figure 5. (A)** *Left,* schematic ofthe experimental strategy. *Right,* WB showing Ser2-RNAPII levels under the different conditions in which the ChIP-seq experiment was performed. DMSO or 100 μM DRB was added to the cells as indicated. **(B)** Genome browser example views of Ser2-RNAPII ChIP-seq data derived from the DRB experiment detailed in *A*. **(C)** *Left,* schematic of the experimental strategy. Briefly,

HeLa WT or Δp300 cells were transfected with siCSB or siCTL 24 h before UV exposure. Then, cells were incubated with DRB and the experiment was performed as in *A*. *Right*, WB showing RNAPII and CSB levels in chromatin-enriched lysate under the different conditions. **(D)** Coding (orange) or template (light blue) strand-biased ChIP-seq signals of the experiment detailed in *C*. **(E)** SSI distribution of HeLa WT (green dots) or Δp300 (blue dots) at 0 min (before DRB wash-out) and 45 min after, with or without UV exposure (20 J/$m^2$), transfected with siCTL or siCSB, as detailed in *C*. All
experiments were performed in at least two independent experimental replicates
