## Supplementary material for "Bypass of Blocking Lesions by RNAPII Impairs the Transcriptional DNA Damage Response": S5

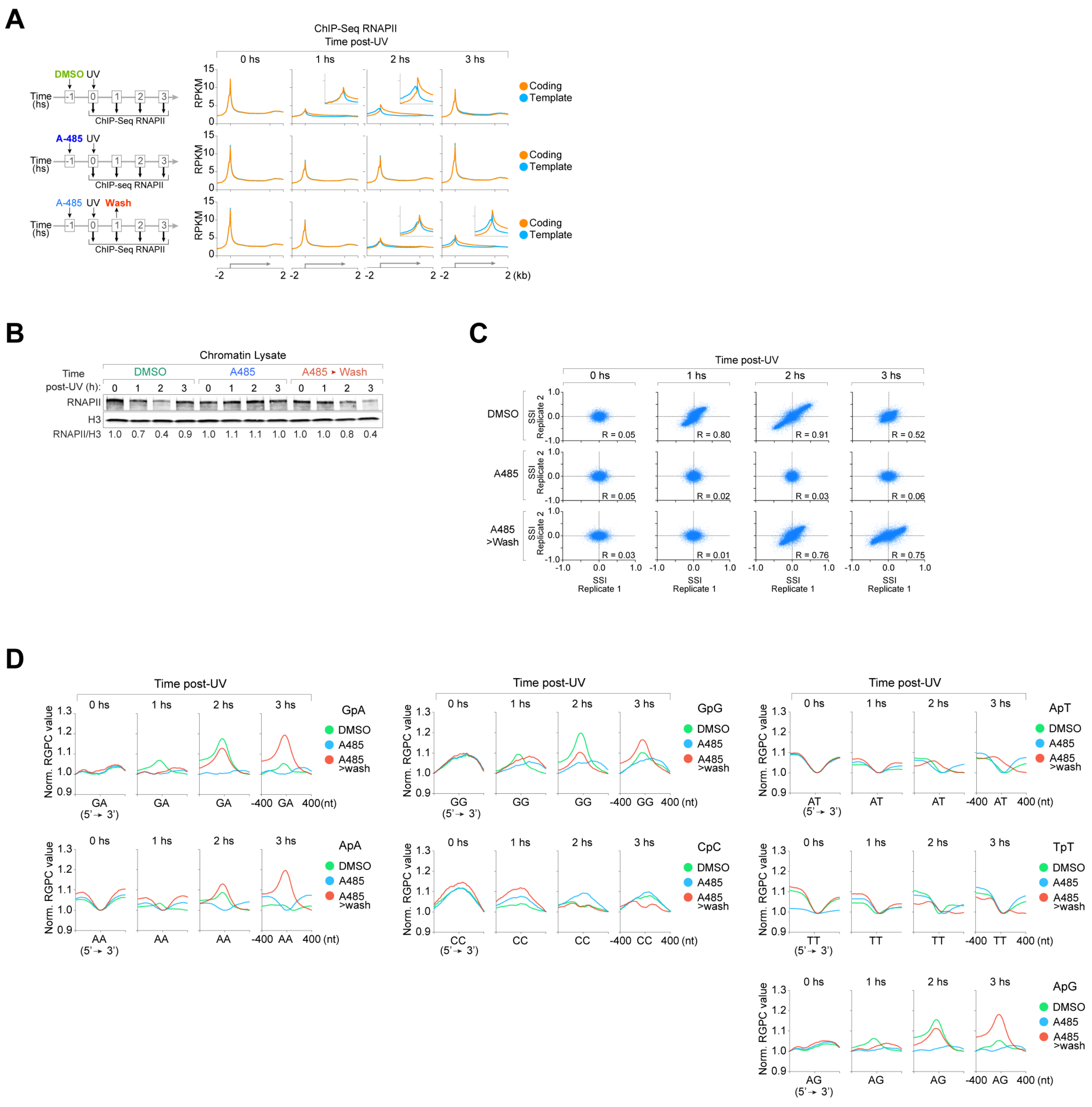


**Figure S5. Related to Figure 5. (A)** *Left*, schematics of the experimental strategy. Briefly, HeLa WT cells were treated with DMSO or 1 μM A485 1 h before UV exposure (20 J/m2). In one group, A485 was washed out 1 h after UV irradiation. Samples were collected at different time points (0 to 3 h after UV exposure). *Right*, metagene analysis showing total RNAPII occupancy on the coding or template DNA strand at the indicated timepoint post UV irradiation (20 J/m$^2$). **(B)** WB analysis of chromatin-enriched fractions of HeLa cells after UV-exposure (20 J/m$^2$). The treatments and time points correspond with those for the ChIP-seq RNAPII experiments detailed in *A*. The ratios of RNAPII and H3 signals relative to DMSO sample before UV damage are shown. **(C)** Correlation of the strand-specific index (SSI) for each gene between experimental replicates in the indicated conditions post UV irradiation. **(D)** Time-course accumulation of RNAPII at different nucleotide dimer sites on the coding strand using the same data as in *Fig. 5C*. Only the central regions of genes longer than 20 kbp were used for the analysis. The lowest value of each profile was used for normalization. All experiments were performed in at least two independent experimental replicates.
