## Supplementary material for "Bypass of Blocking Lesions by RNAPII Impairs the Transcriptional DNA Damage Response": S6

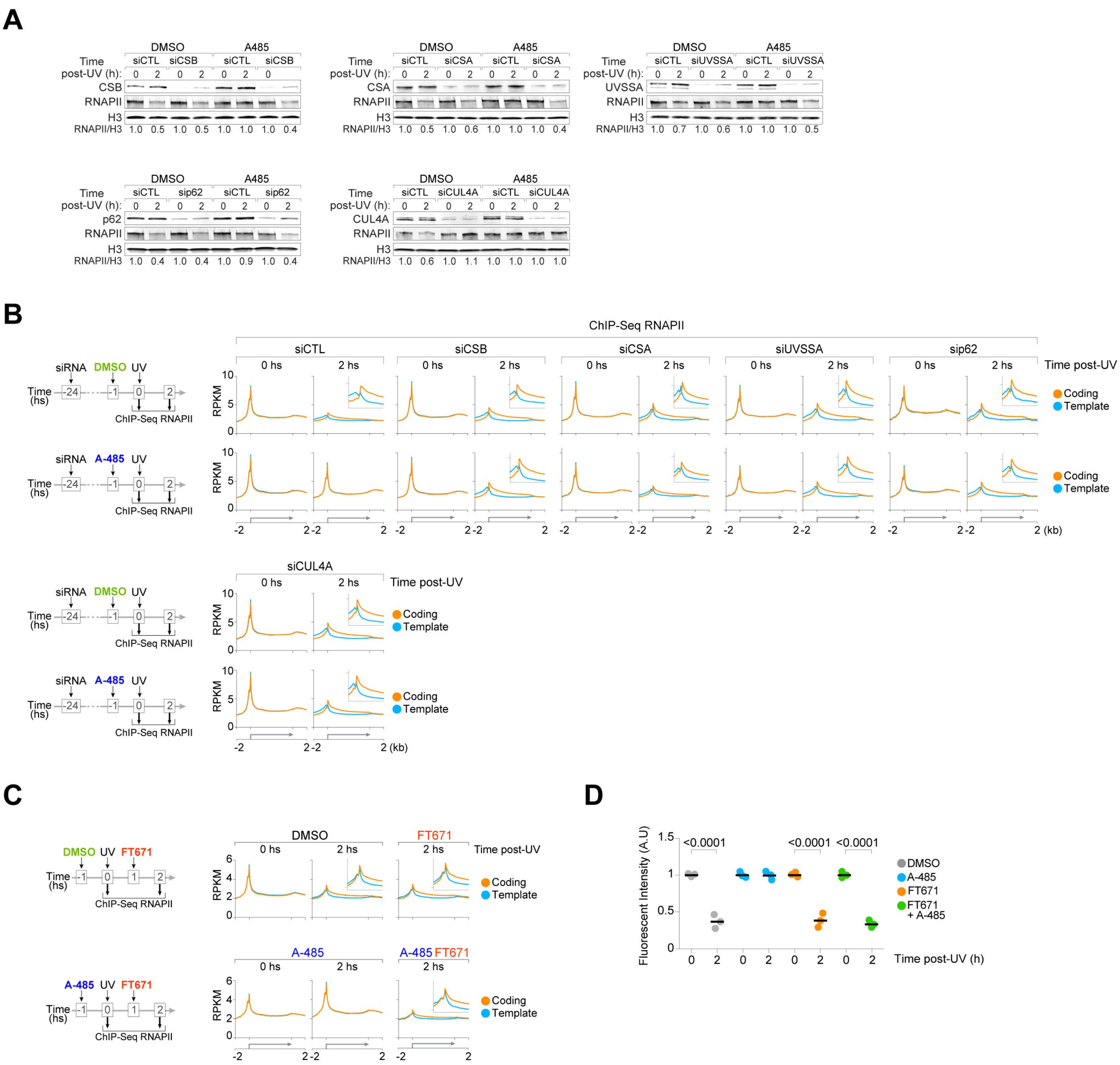


**Figure S6. Related to Figure 5. (A)** WB analysis of chromatin-enriched fractions from HeLa WT cells treated with siRNA (50 nM) of different genes one day before UV exposure (20 J/m$^2$). Cells were incubated with DMSO or 1 μM A485 1 h before UV irradiation. The ratios of RNAPII and H3 signals relative to samples before UV damage are indicated. **(B)** and **(C)** *Left*, schematic of the experimental strategy. *Right*, metagene analysis showing total RNAPII occupancy on the coding (orange) or template (light-blue) DNA strand before and 2h post-UV irradiation (20 J/m$^2$) under the different conditions. **(D)** Quantification of 5-EU signal in HeLa cells at different times after exposure to UV (5 J/m$^2$). Each dot represents the average 5-EU signal per cell, calculated from at least one hundred cells in each experimental replicate (n=3). DMSO or A-485 (1 μM) or FT-671 (1 μM) were added 1 h before UV exposure. 5-EU was added 20 minutes before collection of each sample. P values from one-way ANOVA are shown. All experiments were performed in at least two independent experimental replicates.
