## Supplementary material for "Bypass of Blocking Lesions by RNAPII Impairs the Transcriptional DNA Damage Response": S7

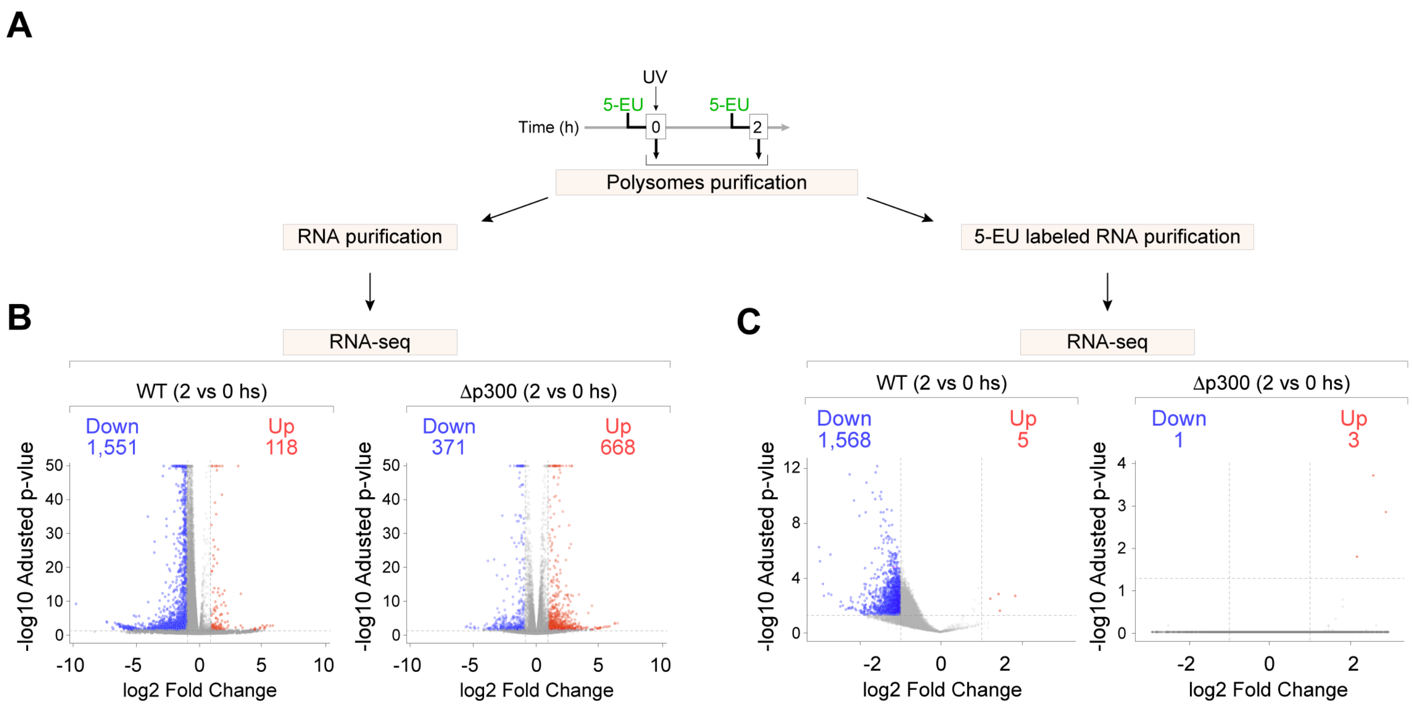


**Figure S7. Polysome analyses reveal that p300-deficient cells sustain production of functional mRNAs after DNA damage. (A)** Schematic of the experimental strategy. **(B)** Volcano plots of differentially expressed genes ($log_2FC \geq 1.5$, q-value < 0.05) from RNA-seq analysis of total polysome-enriched fractions comparing 20 J/m² vs 0 J/m² in HeLa WT or Δp300 cells at 2 h post-UV exposure. Samples were normalized using the *D. melanogaster* spike-in geometric mean. Downregulated genes are shown in blue, and upregulated genes are shown in red. **(C)** Volcano plots of differentially expressed genes ($log_2FC \geq 1.0$, q-value < 0.05) from RNA-seq analysis of 5-EU-labeled RNA purified from polysomal fractions. All experiments were performed in at least two independent experimental replicates.
