## Supplementary material for "Bypass of Blocking Lesions by RNAPII Impairs the Transcriptional DNA Damage Response": S8

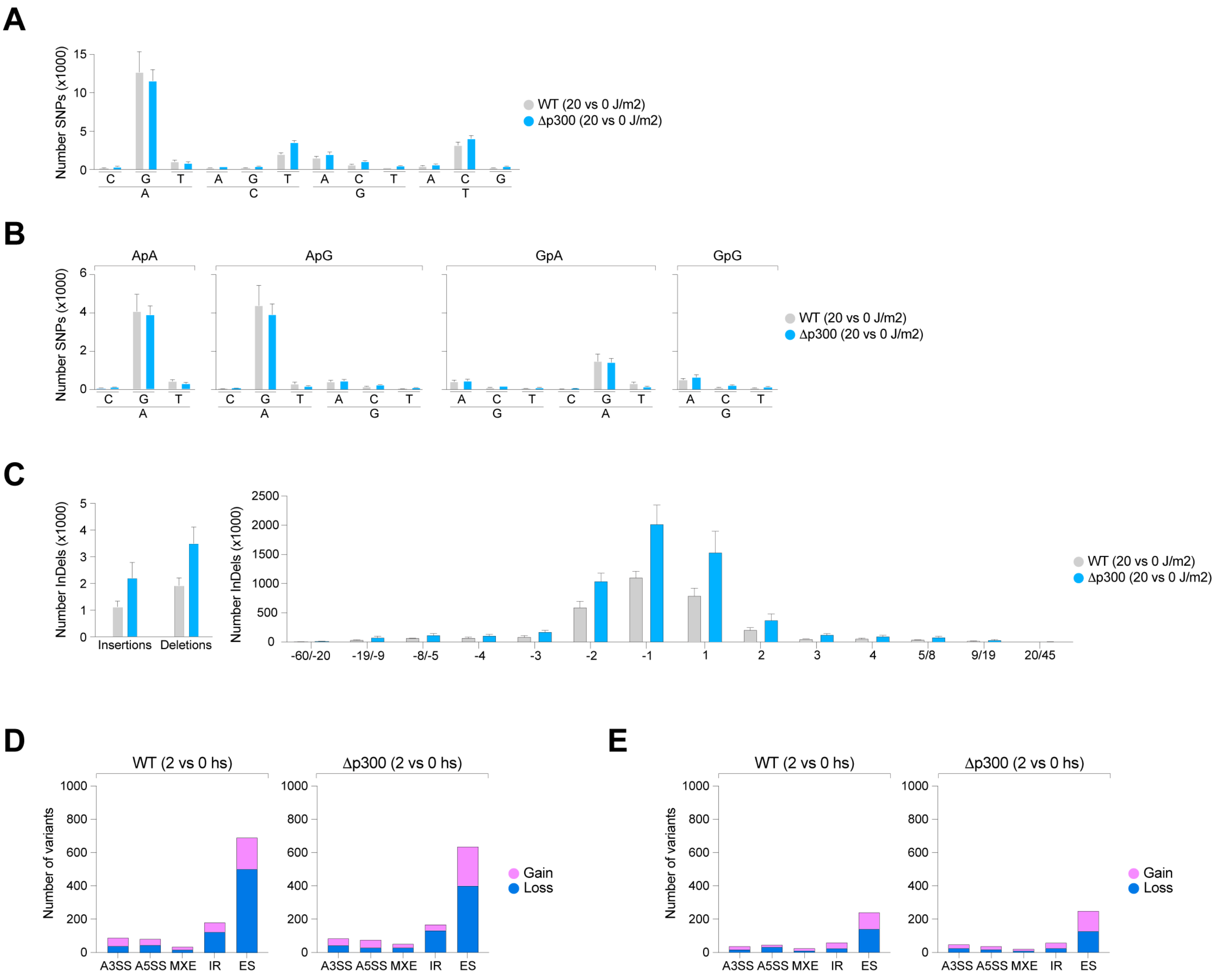


**Fig. S8. P300-deficient cells do not show variation in alternative splicing levels after DNA damage. (A)** Number and types of SNPs exclusively found in ChrRNA-seq of UV-irradiated samples. SNPs also present in control samples were excluded. **(B)** As in *A,* but for SNPs located in different dimer positions on the sense DNA strand of chromosome 1. Only genes oriented in the positive direction were considered for the analysis. **(C)** Insertions and deletions identified exclusively in the ChrRNA-seq of UV-irradiated samples. *Left,* total number of insertions and deletions. *Right,* number of insertions and deletions of varying sizes. **(D)** Splicing analysis of total polysome-enriched fractions RNA-seq of WT (left) or Δp300 (right) HeLa cells. Only variants with ΔPSI ≥ 0.2 or ≤ -0.2, and FDR < 0.05 were considered significant. **(E)** As in *D* but for 5-EU labeled RNA. A3SS: Alternative 3' Splice Site. A5SS: Alternative 5' Splice Site. MXE: Mutually Exclusive Exons. IR: Intron retention. ES: Exon skipping. All experiments were performed in two independent experimental replicates.
