## Supplementary material for "Bypass of Blocking Lesions by RNAPII Impairs the Transcriptional DNA Damage Response": S9

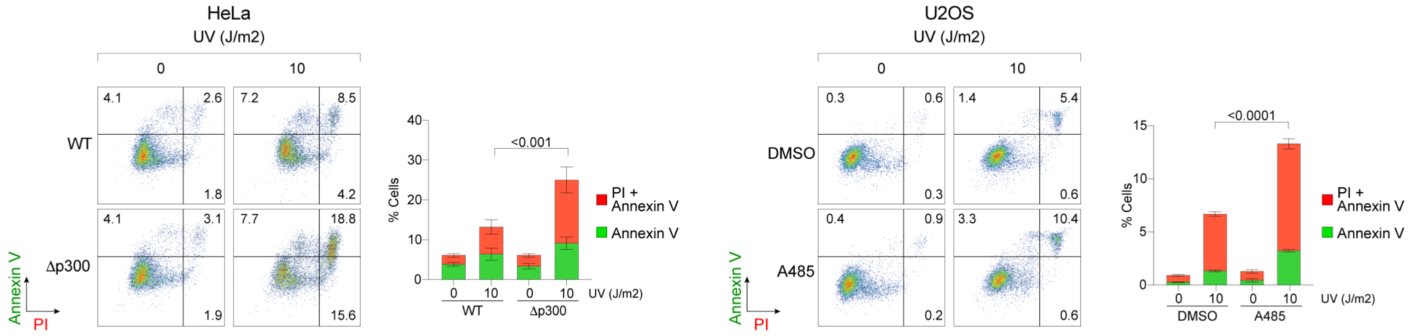


**Figure S9. Related to Figure 6.** Annexin V and propidium iodide (PI) staining in WT and Δp300 HeLa cells (left), and DMSO or 1 μM A-485 treated U2OS cells (right) before and after UV-irradiation (10 J/m$^2$). The percentage of cells positive for only annexin-V (green) or both PI and annexin-V (red) is indicated in bar graphs. All experiments were performed in at least three independent experimental replicates.
